# Reconstruction of the global neural crest gene regulatory network *in vivo*

**DOI:** 10.1101/508473

**Authors:** Ruth M Williams, Ivan Candido-Ferreira, Emmanouela Repapi, Daria Gavriouchkina, Upeka Senanayake, Jelena Telenius, Stephen Taylor, Jim Hughes, Tatjana Sauka-Spengler

**Affiliations:** University of Oxford, MRC Weatherall Institute of Molecular Medicine, Radcliffe Department of Medicine, Oxford, OX3 9DS, UK; University of Oxford, MRC Centre for Computational Biology, MRC Weatherall Institute of Molecular Medicine, Oxford, OX3 9DS, UK; University of Oxford, MRC Molecular Haematology Unit, MRC Weatherall Institute of Molecular Medicine, Oxford, OX3 9DS, UK; Okinawa Institute of Science and Technology, Molecular Genetics Unit, Onna, 904-0495, Japan

**Keywords:** Neural crest, gene regulatory network, chick, enhancers, ATAC, super-enhancers, Capture-C, transcription factor

## Abstract

Precise control of developmental processes is encoded in the genome in the form of gene regulatory networks (GRNs). Such multi-factorial systems are difficult to decode in vertebrates owing to their complex gene hierarchies and transient dynamic molecular interactions. Here we present a genome-wide *in vivo* reconstruction of the GRN underlying development of neural crest (NC), an emblematic embryonic multipotent cell population. By coupling NC-specific epigenomic and single-cell transcriptome profiling with genome/epigenome engineering *in vivo*, we identify multiple regulatory layers governing NC ontogeny, including NC-specific enhancers and super-enhancers, novel *trans*-factors and *cis*-signatures. Assembling the NC regulome has allowed the comprehensive reverse engineering of the NC-GRN at unprecedented resolution. Furthermore, identification and dissection of divergent upstream combinatorial regulatory codes has afforded new insights into opposing gene circuits that define canonical and neural NC fates. Our integrated approach, allowing dissection of cell-type-specific regulatory circuits *in vivo*, has broad implications for GRN discovery and investigation.

## Introduction

Gene regulatory networks (GRNs) are information-processing systems embedded in genomes, responsible for orchestrating cell cycle, homeostasis, physiological processes and development (Levine and Davidson, 2005; Wyrick and Young, 2002). *Cis*-regulatory elements (CREs) form the GRN core, by integrating information from extrinsic signalling cues and endogenous transcription factors (TFs), which together control the spatiotemporal dynamics of complex developmental programmes such as self-renewal and fate decisions (Gouti et al., 2017; Levine and Davidson, 2005; Wyrick and Young, 2002). Pioneering work on GRNs was conducted in several systems, including yeast (Lee et al., 2002), Drosophila (Sandmann et al., 2007), sea urchin embryos (Smith et al., 2007), vertebrate T-lymphocytes (Georgescu et al., 2008) and developing nervous systems (Sandmann et al., 2007; Sauka-Spengler and Bronner-Fraser, 2008), yielding valuable insights into the architecture, logic, modularity and connectivity of developmental circuitries, their disruption in disease, as well as pivotal roles in evolutionary dynamics (Davidson and Erwin, 2006).

GRN dissection has allowed the identification of minimal network topologies such as feed-forward loops and mutually repressive interactions (Davidson, 2010; Milo et al., 2002), which, when combined using powerful algorithmic calculations (for instance through Boolean operations), can accurately predict developmental processes (Peter et al., 2012). However, our understanding of GRNs is still fragmentary, since these insights were implied mainly from studies of transcriptional regulation in unicellular organisms (Lee et al., 2002; Milo et al., 2002) as well as *cis*-regulatory perturbations and epistatic relationships in the sea urchin, Drosophila and vertebrate embryos (Levine and Davidson, 2005; Sauka-Spengler and Bronner-Fraser, 2008), based on a candidate gene approach. Thus, an unbiased genome-wide understanding of vertebrate GRNs has remained elusive.

Vertebrate GRNs use both proximal and distal acting CREs known as transcriptional enhancers (Levine et al., 2014; Levine and Davidson, 2005), that can be located hundreds of kilobases away from their target promoters such as the *Sox9* distal enhancer (Gonen et al., 2018) and within large and often repeat-rich genomes (Prescott et al., 2015), thus posing a challenge for GRN reverse engineering. Nevertheless, single-cell RNA-sequencing (scRNA-seq) recently enabled the interrogation of mouse trunk neural and mesodermal networks (Gouti et al., 2017) as well as those involved in neuronal and cancer circuitries (Aibar et al., 2017), yielding the representation of transcriptome-wide GRNs and new regulatory hierarchies, despite their lack of epigenomic detail. Nonetheless, scRNA-seq coupling to the high-resolution genome-wide mapping of TF binding, chromatin accessibility and histone modifications adapted to low cell numbers allow enhancer characterisation *in vivo* (Georgescu et al., 2008; Smith et al., 2007). Thus, a systems-level approach, combining single-cell transcriptomics and low input epigenomics is poised to resolve complex embryonic GRNs.

The neural crest (NC) is an emblematic embryonic population of multipotent cells that gives rise to a wide range of neural and mesenchymal derivatives including the peripheral nervous system and craniofacial skeleton (Le Douarin and Kalcheim, 1999; Sauka-Spengler and Bronner-Fraser, 2008), thus representing an attractive system to study cell fate decisions and regeneration. Following induction at the neural plate border (NPB) (Basch et al., 2006), premigratory NC cells transiently reside at the dorsal aspect of the neural tube from where they undergo an epithelial-to-mesenchymal transition (EMT), enabling them to migrate, colonise and differentiate within distant sites in the developing embryo (Sauka-Spengler and Bronner-Fraser, 2008), displaying distinct potential and fates at different axial levels (Simoes-Costa and Bronner, 2016). Mistakes and mutations in these processes lead to debilitating birth defects, such as Hirschsprung disease (Carter et al., 2012), craniofacial abnormalities (Trainor, 2010) and CHARGE-syndrome (Schulz et al., 2014), as well as a number of malignancies including melanoma, neuroblastoma and gliomas (Karunasena et al., 2015), highlighting the importance of understanding the global GRN controlling NC ontogeny. Recent mapping of the TFAP2 binding and histone modifications uncovered species-specific enhancer signatures from human and chimp *in vitro* differentiated NC cells involved in craniofacial traits (Prescott et al., 2015; Rada-Iglesias et al., 2012), highlight the power of genome-wide studies in understanding the NC-GRN in development, evolution and disease.

Previous studies have described a number of important NC genes (Simoes-Costa et al., 2014), including evolutionary conserved TFs and downstream effectors leading to a description of a putative NC-GRN underlying key processes during NC ontogeny (Betancur et al., 2010a; Meulemans and Bronner-Fraser, 2004; Sauka-Spengler and Bronner-Fraser, 2008; Simoes-Costa and Bronner, 2015). However, synergistic relationships between components of such GRN were implied from gene-centric knockdown approaches, directionality remained largely unvalidated, and enhancers have only been identified for a small number of factors (Barembaum and Bronner, 2013; Betancur et al., 2010b; Simoes-Costa et al., 2012). Furthermore, other regulatory layers were not integrated into the GRN architecture. For instance, we have recently shown that rather than acting as a canonical transcriptional regulator, FoxD3 specifies premigratory NC programme by priming distal CREs and then arrests NC multipotent state partially through its canonical repressive activity upstream of NC migratory programme (Lukoseviciute et al., 2018). Similarly, Pax7, also known to prime enhancers (Mayran et al., 2018), acts upstream of *Sox10* to drive the premigratory and migratory state and cellular differentiation (Basch et al., 2006; Betancur et al., 2010b; Meulemans and Bronner-Fraser, 2004). However, it has been matter of intense debate whether the NC is a truly homogenous population of multipotent cells or a population of fate-restricted, transcriptionally heterogeneous neural and mesenchymal progenitors (Baggiolini et al., 2015; Bronner-Fraser and Fraser, 1988; Dupin et al., 2018; Le Lievre et al., 1980; Nitzan et al., 2013). Single cell lineage tracing in chick and mice have clearly established multipotency of some NC cells (Baggiolini et al., 2015; Bronner-Fraser and Fraser, 1988) and a stem-cell niche has been proposed (Basch et al., 2006). However, the identification of a sub-circuit in which the EMT factors Snai2 and *Sox9* act upstream of *FoxD3* in the trunk NC suggests a putative role for the latter in the restriction of premigratory NC potential, likely balancing an early restriction of neural and mesenchymal fates (Nitzan et al., 2013). However, NC scarcity, transiency and dynamics in vivo have imposed a considerable challenge to the unbiased interrogation of these conflicting results, and, more broadly, of the complex emergent properties of the developing NC.

Here, we tackle the challenge of reconstructing global NC-GRN by adapting genome-wide epigenome and transcriptome profiling to small numbers of embryonic cells and employing an integrative analysis approach. Using the chicken embryo, we have conducted an unbiased, systems-level functional study of the NC regulome in vivo. By exploring the spatial and temporal dynamics of chromatin accessibility profiles, we identified rewiring of *cis*-regulatory landscapes and dynamic deployment of specific enhancers during NC development. Assigning NC enhancers to downstream target promoters using a high-resolution targeted chromatin conformation capture (3C) based method (Capture-C) (Davies et al., 2016) and positional association to expressed genes, together with the definition of upstream TF inputs using scRNA-seq and motif sequence analysis has enabled comprehensive reverse engineering of the GRN underlying early NC ontogeny. Furthermore, recent advances in genome and epigenome engineering strategies (Williams et al., 2018) have enabled highly efficient, multiplexed knock-out of GRN factors as well as the decommissioning of enhancers providing functional assessment across the GRN.

We have characterised the transcriptional signatures and *cis*-regulatory landscapes intrinsic to subpopulations of NC cells. Our results suggest that within the relatively homogeneous premigratory NC population of multipotent cells that express *Msx2, Alx1* and other stem cell-like factors, the neural (*NR2F2, Pax2, Otx2, Gli2*) and canonical NC identities (*Sox10*, *TFAP2s, Cxcr4, Pax3*) begin to segregate at the onset of delamination. De novo identification of the core transcriptional networks underlying these emergent signatures revealed a combinatorial *cis*-regulatory code comprised of *Sox9*, Otx2, NR2F2 and Pax2 orchestrating NC-derived neural programmes, whereas the canonical regulators *Sox10*, TFAP2, and novel regulators Arnt2 and ATF2 regulate canonical and mesenchymal ones. Through CRISPR/Cas9-mediated knockout of these factors, and their causal link to downstream *cis*-regulatory responsive elements, we reveal direct feed-forward loops controlled by the heterotypic binding of these TFs, as well as the cross-negative regulation of these two identities, which function as core logical features of the NC-GRN. Furthermore, by functional dissection of the *Sox10* super enhancer, we highlight enhancer redundancy and *cis*-regulatory hierarchies as determinants of transcriptional robustness in the NC. Taken together, the ability of our approach to define and interrogate the GRN underlying a system as complex as the NC demonstrates its unique power to dissect gene regulatory circuits *in vivo*, with broad implications for vertebrate GRN discovery and study. Integration of our data sets allowed us to assemble a comprehensive GRN underlying early NC development and provide an interactive resource for exploring new regulatory hierarchies.

## Results

### Transcriptional profiling of early cranial NC

Following electroporation with the cranial NC-specific *FoxD3* enhancer NC1 (Simoes-Costa et al., 2012) driving Citrine (Fig. 1A, B), we used fluorescence-activated cell sorting (FACS) to collect Citrine-positive NC and environing (Citrine-negative) non-NC cells, from developing chicken embryos at two stages: premigratory, 5-6ss (ss, somite-stage) when NC cells maintain a cohesive epithelial structure within the dorsal neural tube where they form a niche of multipotent, self-renewing cells, and early-migratory, 8-10ss, when NC cells have undergone EMT, delaminated from the neural tube and commenced migration.

**Figure 1:**
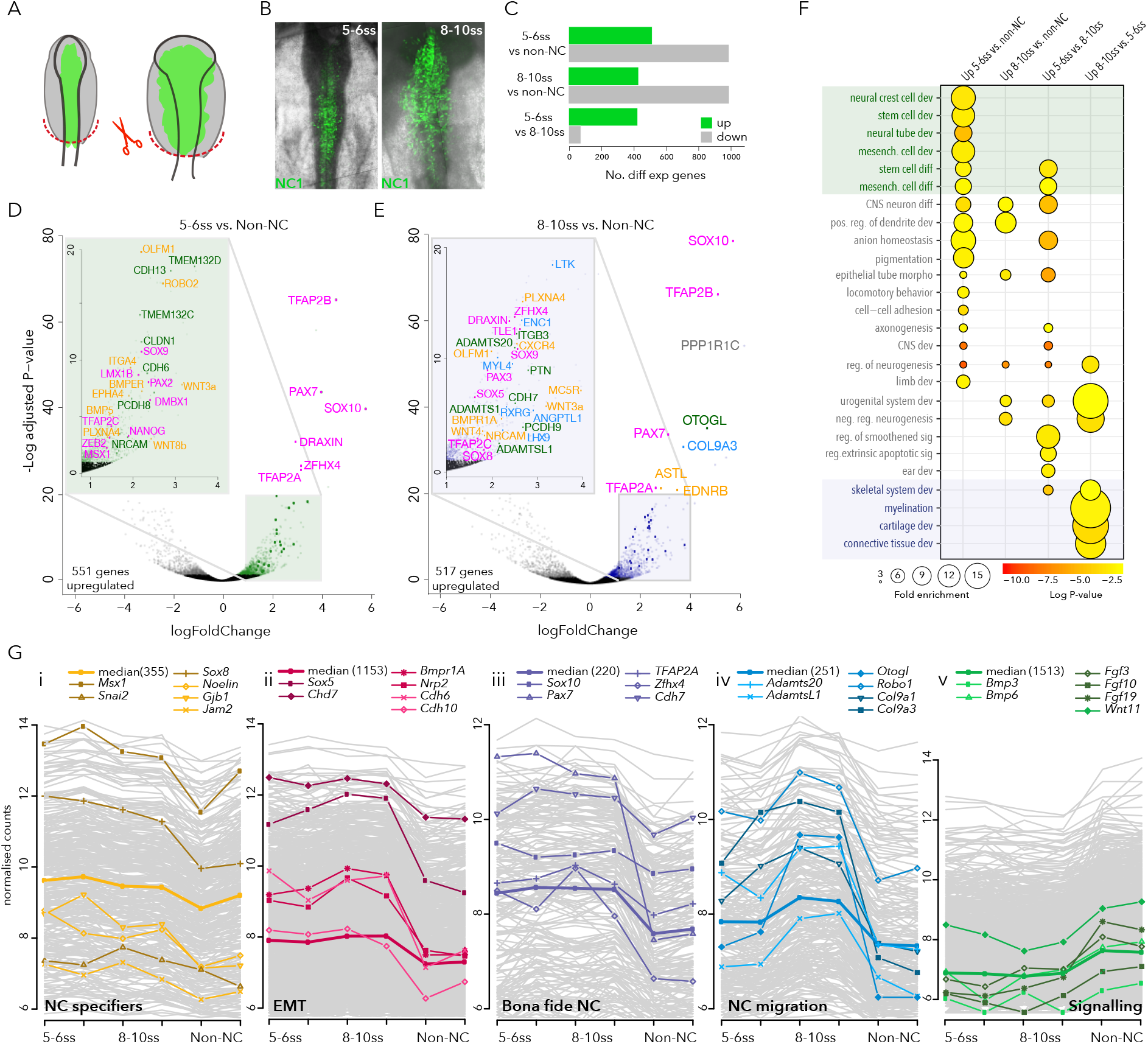
Transcriptional profiling of early cranial NC. **(A)** Schematic representation of Citrine-positive NC cells (green) and Citrine-negative non-NC cells (grey) in early chicken embryos. Red dashed line indicates dissected cranial regions. **(B)** *In vivo* NC specific activity of *FoxD3* enhancer, NC1, at 5-6ss and 8-10ss. **(C)** Number of differentially expressed (enriched and depleted) genes at 5-6ss and 8-10ss compared to non-NC cells, and 5-6ss NC compared to 8-10ss NC. **(D, E)** Volcano plots showing enriched genes (Log Fold Change> 1, base mean>50) at 5-6ss (D) and 8-10ss (E) compared to non-NC cells, magenta; transcription factors, green; cell surface molecules, yellow; signalling molecules, blue; differentiation genes. **(F)** GO terms associated with NC enriched genes. Circle size represents fold enrichment and colour scale denotes the log-scaled statistical significance of enrichment. **(G)** Clusters of highly correlated genes identified by WGCNA, representative genes for each cluster are shown.

RNA-seq on NC (Citrine+) and non-NC (Citrine-) cells followed by differential gene expression analysis revealed approximately 500 genes enriched in NC cells with high reproducibility (Fig.1C; S1A). *Bona fide* NC TFs (*Msx1, Pax7, TFAP2A/B*, *Sox9/10*) and cell adhesion molecules (*Cdh6, Cdh13, Tmem132c/d*) were enriched at 5-6ss, as were signalling components (*Wnt3a/8b, Bmp5, Bmper*) (Fig. 1D). At 8-10ss, *bona fide* NC TFs were maintained and other TFs were added to the programme, such as *Tle1, Sox5/8, TFAP2C, Pax3.* Genes involved in cell migration (*Ednrb, Cxcr4*) and extracellular matrix (ECM) remodelling (*Adamts1/20/l1*) were enriched later as were differentiation genes, reflecting various NC derivative lineages (*Col9a3, Enc1, Ltk, RXRG*) (Fig. 1E; S1B). Consistent with our gene ontology (GO) analysis (Fig. 1F), these results suggest that at 5-6ss, the premigratory NC is engaged in stem cell development and differentiation processes, whereas at 8-10ss they were commencing differentiation programmes (cartilage and connective tissue, myelination and skeletal muscle development). Taken together, we observed a general shift from cell-cell to cell-ECM interactions as NC cells undergo EMT and initiate delamination from the neural tube.

Given the wealth of stages and cell-types transcriptomes, we used Weighted Gene Co-expression Network Analysis (WGCNA) (Langfelder and Horvath, 2008; Langfelder et al., 2008) to further cluster the expressed genes into 13 distinct subgroups, possibly reflecting different modules within the NC programme (Fig.1G and Fig. S1C). Annotation of known NC factors assigns these clusters to previously proposed NC-GRN modules governing critical stages of early NC ontogeny, with further subdivision into early (clusters-i and -ii) and *bona fide* (cluster iii) NC specification modules. NC specifiers (*Snai2, Sox8*) were identified in Cluster-i, along with other early NC regulators *Msx1* and *Noelin1.* Cluster-ii was characterised by NC factors *Sox5* and *Chd7*, as well as cadherins (*Cdh6, Cdh10*) representing the progression of EMT. *Bona fide* NC TFs (*Sox10*, *Pax7, TFAP2A*) were recovered in Cluster-iii and were maintained from 5ss onwards. NC migration was represented by the presence of ECM remodelling factors (*Adamts20, AdamtsL1, Col9a1/a3*) in Cluster-iv. Signalling components required for early NC formation were present in Clusters-v and viii-x where they are predominant in non-NC cells consistent with dynamic signalling interplay of NC cells with environing tissues (Fig.1G and Fig. S1C). Such cluster assignment reflecting co-expression with known NC genes provides an excellent framework from which the role of previously uncharacterised novel factors can be inferred. For instance, we identified *Zfhx4* as a novel NC specifier in Cluster iii, while *Lmo4* and *MEF2A/C*, together with non-canonical Wnt factors (*Wnt7A/B), Plk2* and protein phosphatase 1 regulatory inhibitor subunits (*PPP1Rs*), were recovered in clusters-iv and -vii, the modules harbouring downstream effectors specific to late migration/differentiation into distinct NC lineages.

Overall WGCNA clustering revealed dynamic gene expression modules positively and negatively correlating with NC cells during different stages of their development thus expanding the depth and complexity of the NC-GRN and highlighting the dynamic landscape of the developing NC transcriptome.

### Chromatin dynamics reflects spatiotemporal enhancer activities in vivo

We next set out to identify regulatory features controlling NC programmes. To this end, we performed ATAC-seq (Buenrostro et al., 2013) to generate high-resolution maps of chromatin accessibility, indicative of *cis*-regulatory activity. In addition to NC and non-NC cells at 5-6ss and 8-10ss, ATAC-seq was also performed on dissociated cells from dissected epiblast regions at HH4 (Hamburger and Hamilton, 1951) and somites from HH10 embryos, such that NC could be compared to a more naive and diverse differentiating population, respectively. Linear regression analysis showed a high reproducibility of biological replicates (Fig.S2A, B, C). We observed dynamic changes in chromatin accessibility across the samples analysed. For example, in the vicinity of the NC specifier *Snai2* (Fig. 2A) we identified a number of differentially accessible elements, open specifically in NC cells and not in the naive epiblast, neighbouring tissues or somites (Fig. 2B), we hypothesised these could represent putative NC enhancers. To determine physical interactions between such distal regulatory elements and their putative target genes we performed Next-Generation Capture-C (Davies et al., 2016). Using dorsal neural tubes dissected from 6ss embryos and differentiated blood cells from HH36 embryos as controls we determined the topologically-associating domain (TAD) of *Snai2*, as well as α-globin as control (Fig. 2C). We found a broad NC-specific TAD spanning ~700kb, located predominantly downstream from the promoter and encompassing a number of open chromatin elements. Using *in vivo* reporter assays we identified five active enhancers within the *Snai2* TAD, two distal (enh-332 and enh-334) and three proximal (enh-13, enh-239, enh-242), (Fig. 2D-H; Fig. S3). Enh-13 (Fig.2G-H′) was active in the cranial neural tube including NC at 5ss becoming confined to NC cells by 9ss, representing, to our knowledge, the first NC specific *Snai2* enhancer described to date. Enhancers-332, 334, 242 and 239 were active in both NC and non-NC at 6-10ss (Fig. 2D-F′; Fig. S3), consistent with these regions being accessible in both NC and non-NC cells. A region specifically open in the somites was found to have somite specific activity (enh-241 Fig. S3). Overall these data suggest that chromatin accessibility dynamics across multiple cell types and developmental stages reflect tissue-specific enhancer activity.

**Figure 2:**
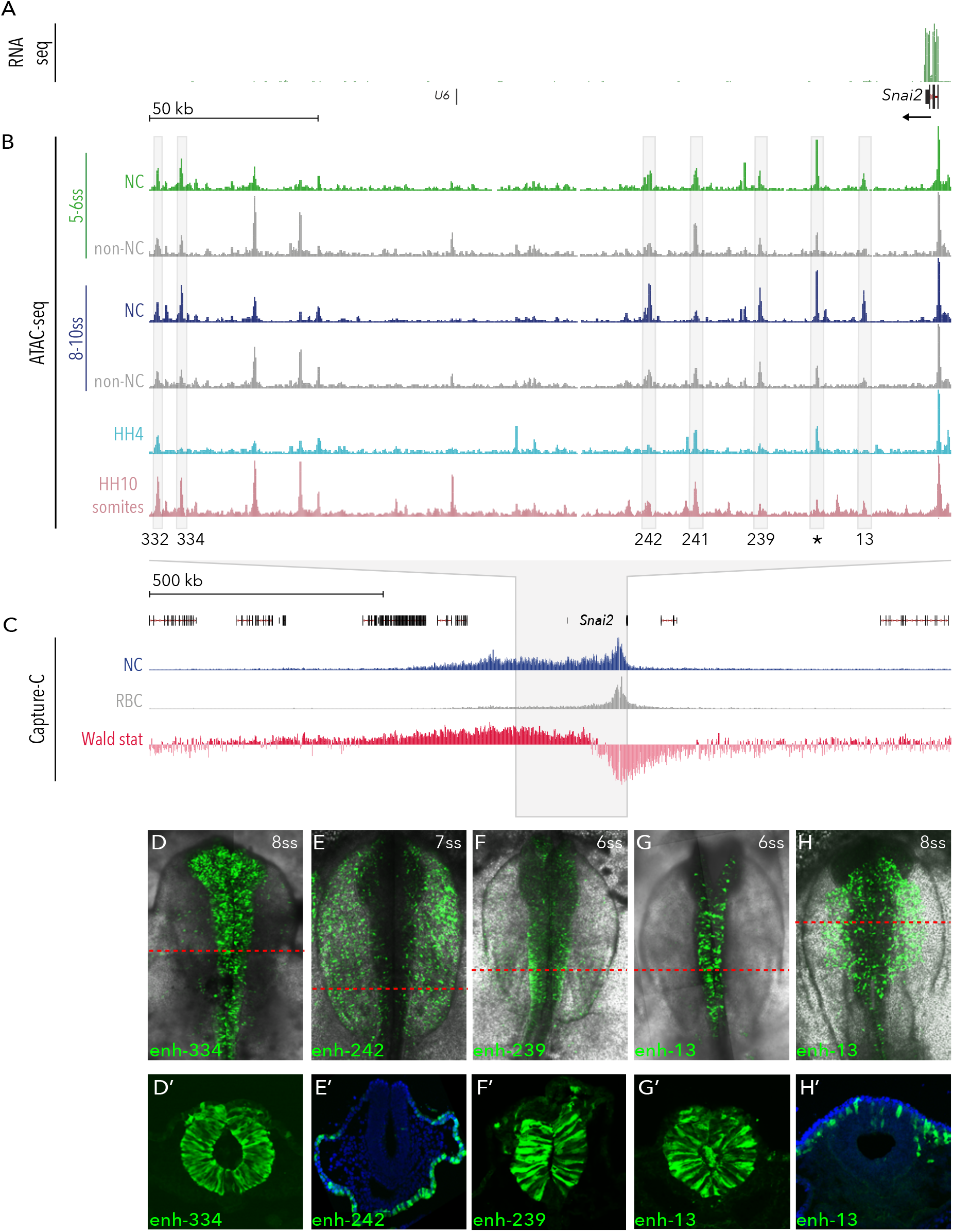
Profiling chromatin accessibility dynamics during early NC development. **(A, B)** Genome browser views of **(A)** 5-6ss NC RNA-seq data and **(B)** ATAC-seq profiles in NC and non-NC cells at the *Snai2* locus, 5-6ss NC ATAC is shown in green and 8-10ss in blue, corresponding non-NC samples are shown underneath in grey. ATAC data from HH4 and somite tissue is shown in light blue and pink respectively. Boxes indicate putative enhancers tested *in vivo.* Asterisk (*) denotes a negative putative enhancer that did not drive fluorescent reporter expression *in vivo.* **(C)** Capture-C tracks showing the TAD at the *Snai2* locus in NC (blue track) and control red blood cells (RBC, grey track). Statistically significant interactions are shown in red track (Wald stat). **(D-H)** Enhancer driven *in vivo* reporter (Citrine) expression of tested enhancers. (**D**–*H*) Transverse sections of (D-H), immunostained for Citrine, red line indicates approximate location of section, blue, DAPI. Read counts were normalised and scale bars are consistent across all samples shown.

### Dynamics of *cis*-regulatory landscapes

To expand the analysis of chromatin dynamics globally and attain the regulatory information encoded in enhancers, we performed *k*-means clustering of our ATAC-seq datasets. Searching for patterns of differential chromatin accessibility genome-wide, we identified ten distinct clusters of elements, and we focused on the top seven clusters based on their density and patterns of accessibility (Fig. 3A and Fig. S2E). Broad signal elements constitutively accessible across all samples were recovered in *k*-Cluster-4 (Spearman correlation coefficient rS=0.94) (Fig. 3A, B); 20% of which represented promoter regions (Fig. S2D). Elements specifically accessible in NC cells were found in *k*-Cluster-3 (r_S_=0.74, *p*<2.2 x 10^−16^) (Fig. 3A, B), which we assigned as the *bona fide* NC enhancer group. Accessibility in NC cells also characterised the *k*-Cluster-9 elements (r_S_=0.74, *p<2.2* x 10^−16^) (Fig. 3A, B); however, these were also open in Citrine-negative non-NC cells, indicating possible activity in the environing neuroepithelial ectoderm. Chromatin accessibility was also seen in premigratory NC and in the early (HH4) epiblast (*k*-Cluster-6), consistent with previous reports suggesting that the NC regulatory landscape is established as early as during gastrulation (Basch et al., 2006; Buitrago-Delgado et al., 2015). *K*-cluster-6 accessibility was reduced in NC cells at 8-10ss, suggesting these elements were involved in the early establishment of NC programmes and later decommissioned (Fig. 3A). Elements more prominently open in non-NC but maintaining some accessibility in NC were recovered in *k*-Cluster-2. Early (HH4) acting elements were found in *k*-Clusters −5 and −7. *K*-Cluster-7 elements were more dynamic; compacted in NC cells at 5-6ss, but open in non-NC at the same stage, and open in both NC and non-NC at 8-10ss, possibly regulating late neural fates (Fig. 3A). Elements recovered in dynamic *k*-clusters were predominantly located in intergenic or intronic regions (Fig. S2D), consistent with their *cis*-regulatory activity and were found to correlate with unimodal transcription (Fig. S2F, F’). These results thus establish the *cis*-regulatory landscape dynamics across different stages and cell-types, likely underlying NC ontogeny.

**Figure 3:**
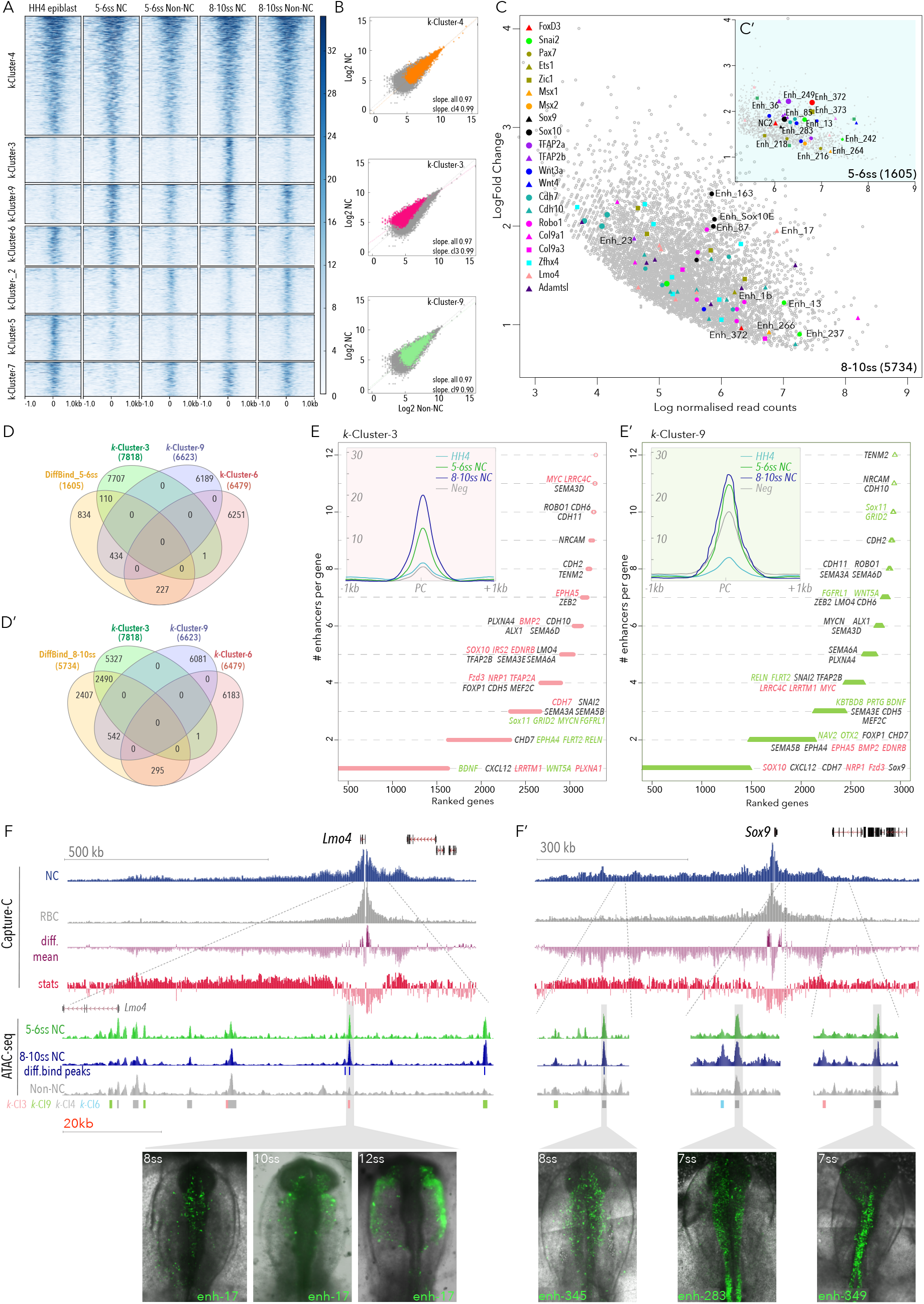
*K*-means clustering and differential analysis of chromatin accessibility during early NC development. **(A)** Heatmap depicting *k-*means linear enrichment clustering of ATAC signal across stages and cells analysed. Corresponding correlation plots for selected clusters are shown in **(B)**. **(C)** MA plot depicting differentially accessible elements (DiffBind). A selection of putative enhancers to key NC genes are colour coded, numbered enhancers have been tested (Fig. 2D-H, Fig. 3F, F′, Fig. 4C-H and S3). **(D, D′)** Venn diagrams demonstrate overlap of *k*-Cluster and DiffBind elements, total number of elements in each is shown. **(E, E′)** Plots representing genes assigned to *k*-Cluster-3 (E) and *k*-Cluster-9 (E′) elements, ranked by the number of associated elements. Inserts show mean merged profiles of accessibility across all samples. **(F, F′)** Genome browser snapshots of *Lmo4* (F) and *Sox9* (F′) loci. Showing Capture-C (blue, NC cells, grey, red blood cells (RBC), purple, differential mean between experimental and control samples, red, statistically significant interactions)) and ATAC (green, 5-6ss, blue, 8-10ss, grey, negative non-NC) tracks. *k*-means elements are also shown. Selected putative enhancers are boxed in grey and their activity is shown in underlying embryos images.

In parallel, we employed a stringent statistical analysis of differential accessibility in purified NC cells compared to neighbouring non-NC cells using the DEseq2 algorithm, part of the DiffBind package (Stark, 2011). We identified 1605 statistically significant NC-specific elements (from now on referred to as DiffBind elements) at 5-6ss and 5734 at 8-10ss (FDR<0.1, Fold change>1) (Fig. 3C, C′), suggesting the progressive establishment of the *bona fide cis*-regulatory landscape during NC development. Diffbind elements were enriched at key NC loci (Fig. 3C, C′) and they differentially overlapped individual *k*-Clusters. Consistent with their early opening, DiffBind elements at 5-6ss were mainly shared with *k*-Cluster-9 (~30%), *k*-Cluster-6 (~14%), and to a lesser extent with *k*-Cluster-3 (~7%). DiffBind 8-10ss elements predominantly overlapped with *k*-Cluster-3 elements (~45%), associated with bona fide NC genes, and to a lesser degree with *k*-Cluster-9 (~10%) and *k*-Cluster-6 (~5%) (Fig. 3D, D′).

Assignment of *cis*-regulatory elements from *k*-Clusters −3 and −9 to the nearest expressed gene indicated that key developmental NC genes are controlled by multiple regulatory elements. We further distinguished genes predominantly regulated either by specific *k*-clusters and those with shared regulation from two or more *k*-clusters. For example, canonical NC factors *Sox10*, *Ednrb* and *Myc*, along with lesser-characterised NC genes such as *Lrrc4c* and *Epha5* were enriched for *k*-Cluster-3 elements (Fig. 3E). In contrast, genes involved in neural development such as *Sox11, Grid2, MycN* and cell migration/guidance *Fgfrl1, Wnt5a* and *Sema3a* were largely enriched for *k*-Cluster-9 elements (Fig. 3E′). However, other *bona fide* NC genes *TFAP2B* and *Snai2*, and EMT factors *Zeb2* and *Lmo4* were driven by a combination of *k*-Cluster-3 and *k*-Cluster-9 elements (Fig. 3E, E′), suggesting a complex interplay between cell-types, developmental stages and *cis*-regulatory dynamics. Collectively these results suggest that distinct clusters of enhancers may differentially regulate different NC lineages.

We next validated genome-wide enhancer predictions by assessing the spatial and temporal activity of an extensive cohort of putative enhancers *in vivo.* We selected a number of canonical *bona fide* NC loci (*Sox10, Pax7, Sox9, Snai2, Ets1, TFAP2A* and *TFAP2B*), as well as *Lmo4*, and determined their TADs by Capture-C. TADs varied in size and distribution, for example, the *Lmo4* TAD spread ~500kb downstream and ~200kb upstream of the promoter (Fig. 3F) whereas the *Sox9* TAD spanned evenly across a large (~1.5Mbp) region up- and downstream from the promoter (Fig. 3F′). The ~750kb *TFAP2B* TAD spread predominantly downstream (Fig. S4A) whereas the *Pax7* TAD (~500kb) predominantly encompassed the body of the *Pax7* gene and included mostly intronic and downstream *cis*-regulatory elements (Fig. S4B). We developed a novel high-throughput screening assay using a modified highly-efficient GoldenGate cloning strategy combining fluorescent and ′Nanotag′ reporters (Nam and Davidson, 2012) that are also detectable by Nanostring (see methods and Fig. S3A-C). We tested the activity of a number of novel enhancers regulating NC genes including *Lmo4* (enh-17) (Fig. 3F), *Sox9* (enhs-283, −345, −249) (Fig. 3F′), *Pax7* (enhs-194, −195, −199, −216, −218), *FoxD3, TFAP2A* and many others (Fig. S3D). These results validate our integrative framework for identification of cell-type- and stage-specific NC enhancers and provide an excellent resource for the study of NC gene regulation.

Taken together, our results suggest that differential chromatin accessibility analysis and *k*-means clustering successfully captured *cis*-regulatory landscape dynamics, providing an insight into the enhancer usage and deployment during NC ontogeny, as well as into the nature of interactions shared with other cell types. We propose that *k*-Cluster-6 elements initiate NC specification, likely through *cis*-regulatory pleiotropy, for example, reusing enhancer elements from epiblast cells in the premigratory NC. Establishment of an early *bona fide* NC programme is captured by *k*-Cluster-9 elements, consistent with the NC origin at the dorsal neuroepithelium. We, therefore, hypothesise that *k*-Cluster-9 might represent NC and non-NC neural progenitors, to which the early, premigratory NC share regulatory circuits. In contrast, *k*-Cluster-3 elements may represent spatially restricted, *bona fide* NC. DiffBind clusters at 5-6ss and 8-10ss capture the temporal dynamics of canonical premigratory and early delaminating *bona fide* NC, revealing a global picture of NC spatiotemporal heterogeneity at the epigenomic level. Thus, we hypothesised that the regulatory logic of NC development is hierarchically embedded in the dynamic *cis*-regulatory landscapes. This approach has thus allowed us to infer the full complement of active NC enhancers, from which the NC-GRN could be reconstructed.

### Enhancer hierarchies within a super-enhancer dynamically controlling *Sox10*

The observed spatial organisation of dynamically accessible regions (Fig. 3F, F′, Fig. S4), as well as the increased ratio of enhancers per gene at developmental loci (Fig. 3D, D′, S2G) suggests that enhancers form clusters of elements within TADs, constituting locus control regions (Spitz et al., 2003; Talbot et al., 1989) also known as super-enhancers (SEs) (Hay et al., 2016; Hnisz et al., 2013). Importantly, SE-like regions function as the core regulatory circuits within GRNs, specifying the cell-type identity and their dysregulation can impair cellular programming and be involved in the emergence of neoplasias (Boeva et al., 2017; Hay et al., 2016; Hnisz et al., 2013). Given their functional importance, we have performed ChIP-seq in NC cells for H3K27Ac, as SE-like regions are differentially enriched for this histone modification (Loven et al., 2013). Using the ROSE (Rank Ordering of Super Enhancers) algorithm (Loven et al., 2013; Whyte et al., 2013) on H3K27Ac ChIP-seq profiles, we identified 1379 SEs (Fig. 4A) at the delaminating NC stage (8-10ss), and 1288 SEs at the premigratory NC stage (5-6ss) (Fig. S5A). Top-ranked SE-like clusters were associated to canonical NC regulators loci, including *Sox10*, *Fzd3, TFAP2A, TFAP2B, Sox5* and *Snai2* (Fig. 4A) at 8-10ss, but not at the earlier premigratory stage, suggesting rewiring of the *bona fide* NC *cis*-regulatory landscape at delaminating/actively migrating stage, when the majority of enhancers are deployed. Moreover, occupancy of the Mediator complex as inferred by ChIP-seq of the transcription co-factor Brd4 suggests that approximately 30% of *bona fide* NC elements from the Diffbind 8-10ss cluster belong to SE-like regions (Fig. 4B).

**Figure 4:**
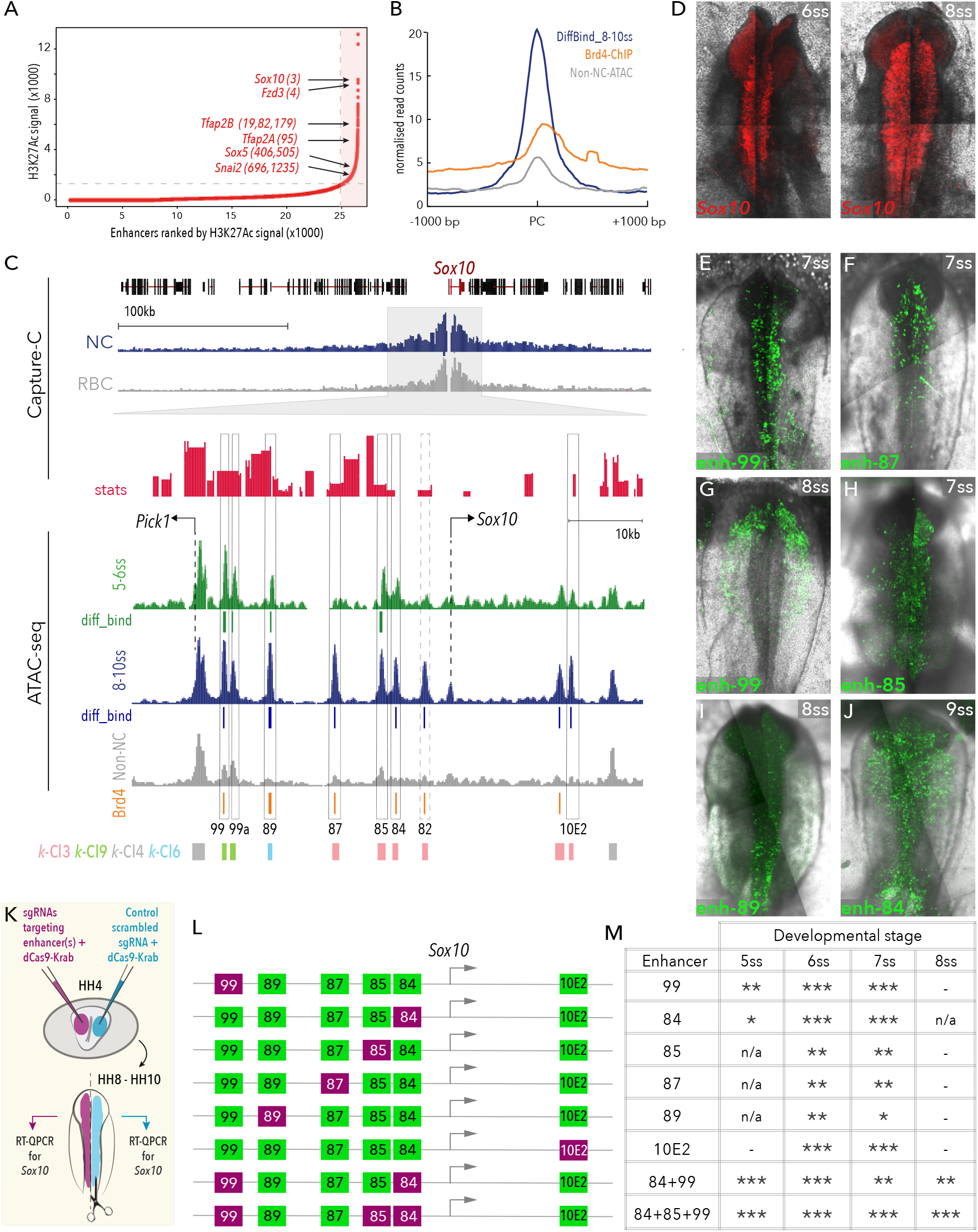
Super-enhancer-like clusters regulate key NC genes. **(A)** Enhancers ranked by H3K27ac signal from 8-10ss NC, using the ROSE algorithm, top-ranked NC genes are annotated. **(B)** Mean merged profile of DiffBind elements at 8-10ss and control non-NC elements occupied by Brd4. **(C)** Genome browser tracks of Capture-C (blue, NC cells, grey, RBC, red, statistically significant interactions) and ATAC data (green, 5-6ss, blue, 8-10ss, grey non-NC) at *Sox10* locus. DiffBind elements 5-6ss (green) and 8-10ss (blue), Brd4 called peaks (orange) and *k*-means elements are shown under ATAC tracks. Selected putative enhancers are boxed. **(D)** Endogenous *Sox10* expression pattern detected by fluorescent *in situ* (hybridisation chain reaction, HCR). (E-J) *in vivo* activity of novel *Sox10* enhancers. (K) Schematic of bilateral electroporation assay for epigenome engineering experiments. (L) Schematic representation of enhancers targeted for decommissioning. (M) Table describing effect on endogenous *Sox10* expression following CRISPR mediated epigenome modification of *Sox10* enhancers (* weak effect (<20% decrease in *Sox10* expression); ** moderate effect (20-40% decrease); *** strong effect (>40% decrease); n/a not analysed; - no effect).

Given the pivotal role of *Sox10* in NC ontogeny (Kelsh, 2006; Sauka-Spengler and Bronner-Fraser, 2008) and its top SE ranking (Fig. 4A), we sought to thoroughly survey the *Sox10* locus at the epigenomic level. We identified a cluster of open elements spanning a ~20kb region, 2kb upstream from the promoter, within the defined *Sox10* TAD (Fig. 4C). Each of these elements were present in our DiffBind sets and the majority were recovered in *k*-Cluster-3, with the exception of the two most distal peaks (enh-99 and enh-89), found in *k*-Cluster-9 and *k*-Cluster-6, respectively (Fig. 4C), consistent with their accessibility dynamics. *In vivo* enhancer reporter assays showed that these elements collectively reconstituted the endogenous *Sox10* expression (Fig. 4D-J), albeit with differing spatiotemporal patterns of activity. Enh-99 identified in *k*-Cluster-9 and co-occupied by Brd4, is the earliest acting *Sox10* enhancer (detected from 5ss) that likely onsets the *Sox10* expression (Fig. 4E, G). Interestingly, we found a region of Enh-99 (Enh-99a) predominantly active in trunk NC cells (Fig. S3D). This is followed by enh-87, −85, and −84 which onset at 7ss (Fig. 4F, H) and continue to be active throughout early NC migration, with enh-84 activity particularly prominent in migrating NC (8-10ss) (Fig. 4J). This is consistent with the occupancy of these elements by Brd4 (with the exception of enh-85) and their assignment to the *bona fide k*-Cluster-3. Surprisingly, Enh-89, a *k*-Cluster-6 element accessible from the epiblast stage, is active at 8ss, predominantly in the hindbrain (Fig. 4I), providing further evidence of the role of enhancer pleiotropy (Preger-Ben Noon et al., 2018) in NC gene regulation. Conversely, enh-87, a *k*-Cluster-3 element, is active in a complimentary domain (anterior cranial region, Fig. 4F), suggesting that enh-89 and enh-87 collectively control cranial antero-posterior *Sox10* expression in a manner redundant to enh-99, possibly acting as shadow enhancers (Cannavo et al., 2016). All five new enhancers displayed statistically significant interactions with the *Sox10* promoter (Fig. 4C). The most proximal open element, enh-82, which did not display any *in vivo* enhancer activity at the stages tested, possibly represents a *cis*-regulatory repressor or a late-acting enhancer. Interestingly, although both previously described *Sox10* enhancers (Betancur et al., 2010b) were recovered by our DiffBind analysis, only cranial enhancer 10E2 and not the trunk enhancer 10E1, showed significant interaction with the *Sox10* promoter, suggesting an instructive model (de Laat and Duboule, 2013) in which *de novo* formed loops control *Sox10* transcription at different axial levels.

To determine the functional contribution of each discrete element within this SE-like region to *Sox10* transcription we used CRISPR-mediated epigenome engineering to modulate endogenous enhancer activity (Williams et al., 2018), whereby five guide RNAs (gRNAs) were employed to target the transcriptional repressor dCas9-Krab to each individual enhancer. We used an *in vivo* bilateral electroporation assay, in which the left side of the embryo received targeted gRNAs and dCas9-Krab and the right side received a scrambled control gRNA with dCas9-Krab (Fig. 4K). Left and right dorsal neural tubes from individual embryos were dissected at 5-8ss and QPCR was performed to determine the effect of enhancer perturbation on endogenous *Sox10* expression (Fig. S5B). We found that loss of enh-99 alone affected *Sox10* expression early (5ss), whereas targeting other enhancers only disrupted *Sox10* expression from 6-7ss (Fig. 4L, M). However, decommissioning any individual enhancer on its own was insufficient to cause a lasting effect on *Sox10* transcription (past 7ss), probably due to compensation by other concurrently active elements. *Sox10* expression indeed increases as NC development progresses, and such enhancement is most likely due to multi-enhancer usage. We thus next targeted other *Sox10* enhancers in conjunction with enh-99 to determine their contribution to increase and maintain *Sox10* transcription. We found that simultaneous repression of enh-84 and enh-99 was sufficient to disrupt *Sox10* expression from 5ss through to 8ss (Fig.4L, M; Fig. S5B). Furthermore, when enh-84, −85 and −99 were decommissioned concurrently, *Sox10* transcription was continually repressed through to the 8ss (Fig. 4L, M; Fig. S5B). Taken together, these results suggest that additive or sub-additive (Bothma et al., 2015; Hay et al., 2016), combinatorial action of enhancers is critical for sustained *Sox10* expression during NC EMT and demonstrates functional redundancy of individual enhancers as previously described in other systems (Osterwalder et al., 2018). Moreover, our results provide a locus specific example of the global dynamics of NC transcriptional regulation at a key SE, by providing evidence that: (1) *k*-Cluster-9 elements may be required and sufficient for initiation, but not for maintenance of NC-specific programme, as in the case of enh-99, and are shared with neural progenitor cells (such as trunk neural tube cells, Enh-99a); (2) early regulatory elements from stem cells are (re)used in NC cells (Buitrago-Delgado et al., 2015), as revealed by *k*-Cluster-6 member enh-89; and (3) multiple *k*-Cluster-3 elements (enhs-84, 85, 87 and 10E2) represent *bona fide* NC enhancers essential for transcriptional robustness within the NC-GRN.

### Early heterogeneity and NC lineage decisions

The observed early *cis*-regulatory dynamics (Fig. 3, 4, Fig. S4) provides evidence of early heterogeneity at the epigenomic level, consistent with previous reports suggesting regulatory mechanisms that limit NC potential and thus determine NC lineage decisions, with the implications for the proposed multipotency of the NC (Dupin et al., 2018; Nitzan et al., 2013). Consistently, GO term enrichment analysis of genes associated to each group of clustered elements revealed that developmental pathways associated with canonical NC development and differentiation programmes (*p*<0.01, Binomial test with Bonferroni correction) were exclusively associated with *k*-Cluster-3 and DiffBind elements (Fig. 5A, red box). *k*-Cluster-3 and DiffBind enhancers only become accessible from premigratory NC stages (Fig. 5B), providing further evidence that they represent linchpin elements mediating *bona fide* NC programmes. Conversely, programmes associated with neural lineages are shared (and hierarchically regulated) by the successive action of *k*-Cluster-6, −9 and −3 elements (Fig. 5A, green box), revealing an early regulatory split between mesenchymal and neural progenitors at the *cis*-regulatory level. Nevertheless, DiffBind elements at 5-6ss suggest the existence of a stem cell-like niche within *bona fide*, premigratory NC, likely maintained at least until 8-10ss (Fig. 5A, red box).

**Figure 5:**
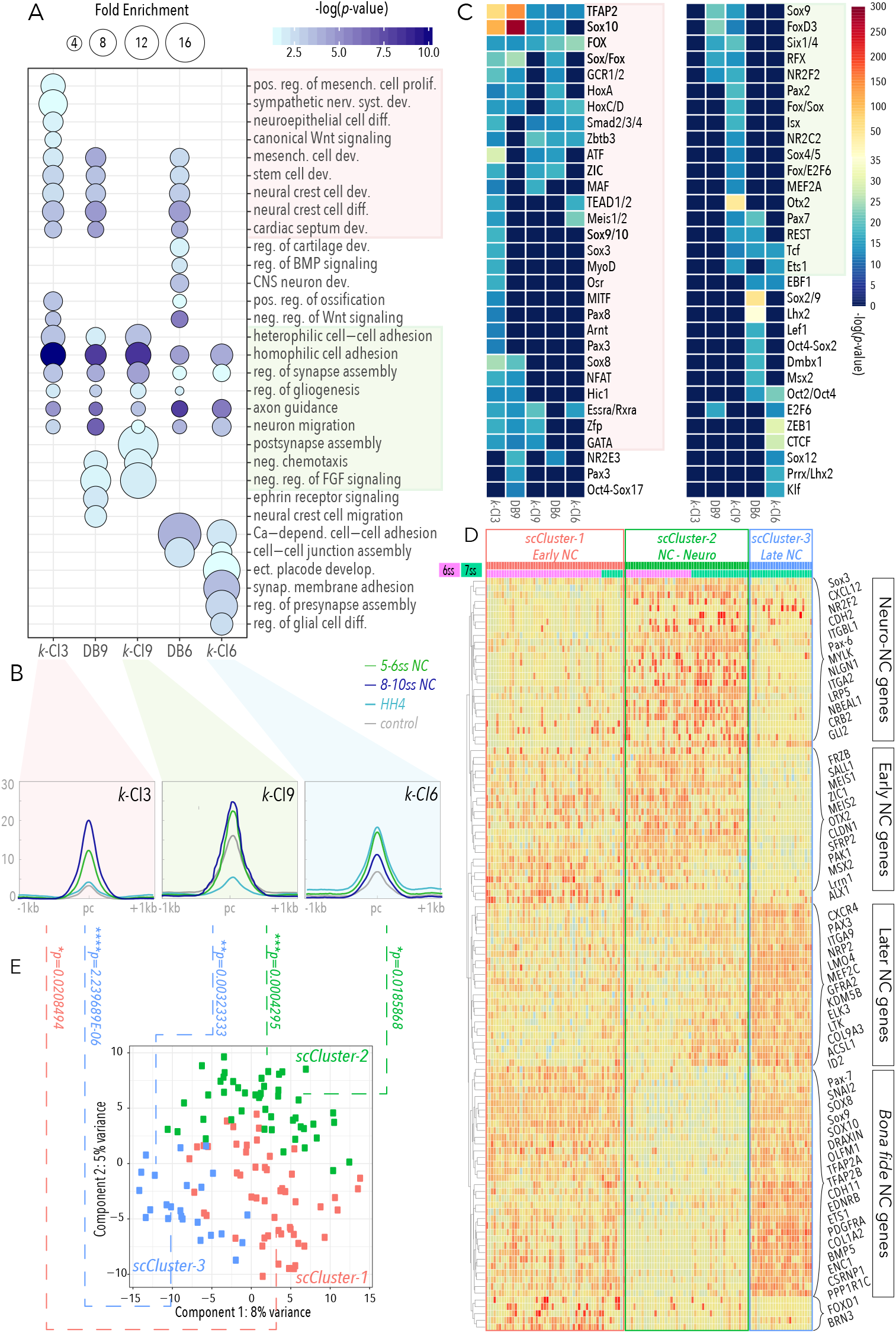
Functional dissection of *k*-Cluster elements and assignment to single-cell transcriptomes. **(A)** GO terms yielded from *k*-Cluster and DiffBind associated genes. Circle size represents fold enrichment and colour scale denotes the log-scaled statistical significance of enrichment. *K*-Cluster-3 and DiffBind elements exclusively associate with GO terms reflecting canonical NC and mesenchymal cell development, and specific NC differentiation programmes (sympathetic nervous system), while sharing roles in homophilic cell adhesion, gliogenesis and axonogenesis with *k*-Clusters −6 and −9 (**p<0.01, Binomial test with Bonferroni correction, fold change>4). Only late-acting elements (*k*-Cluster-3, *k*-Cluster-9 and DiffBind 8-10ss) correlate to heterophilic cell-cell adhesion. *K*-Cluster-6 elements associate with early regulation of neuronal NC lineages as well as with ectodermal placode and cell-cell adhesion phenotypes. **(B)** Mean merged density profiles of *k*-Clusters-3, −9 and −6. *K*-Cluster-3 elements begin opening at 5-6ss when *k*-Cluster-6-9 are already open, at 8-10ss *k*-Cluster-3-9 are fully accessible and *k*-Cluster-6 are elements are becoming compressed following exclusive accessibility at HH4. *K*-Cluster-3-6 elements show limited accessibility in non-NC, whereas *k*-Cluster-9 show modest opening in non-NC. **(C)** *De novo* TF binding motifs enriched in *k*-Cluster and DiffBind elements. **(D)** Single-cell-RNA-seq heatmap visualising hierarchical clustering of single NC cells at 6-7ss (total 124 cells, 74 6ss and 63 7ss). Top 50 differentially expressed genes are shown. **(E)** PCA of top 100 genes from sc-RNA-seq analysis. sc-cluster-1 genes (early NC, shown in red) associate to *k*-Cluster-3 elements with statistical significance (*p*=0.021), sc-cluster-3 genes (late NC, blue) associate to *k*-Cluster-3-9 (*p*=2.24×10^6^, p=0.003 respectively) and sc-cluster-1 (neuronal-NC, green) associate with *k*-Cluster-6-9 (*p*=0.018, p=0.0004 respectively).

In addition to sharing NC terms with *k*-Cluster-3 and *k*-Cluster-9, DiffBind elements yielded terms indicative of the current identity (early multipotent, premigratory NC at 5-6ss, and early differentiating, migratory NC at 8-10ss), as well as highlighting the future state of NC cells relative to the sample stage. For instance, terms including cartilage development, regulation of ossification and CNS neuron development, more reflective of later NC behaviour, were over-represented at 5-6ss, whereas terms such as gliogenesis and neuron migration were enriched at 8-10ss, suggesting that the *cis*-regulatory landscapes are primed earlier than the onset of corresponding gene expression and observable phenotype. Interestingly, early-opening *k*-Cluster-6 elements (Fig. 5B) function upstream, in the regulation of neuronal differentiation processes (pre-synapse assembly, synaptic membrane adhesion and gliogenesis), suggesting that at the regulatory level, neural-NC fates begin to be primed from as early as gastrulation. This is in line with the finding that all *k*-Cluster-9 elements that associate with neural differentiation appear to be fully accessible by 5-6ss (Fig. 5B).

We next investigated the upstream *cis*-regulatory codes driving the observed enhancer heterogeneity using the Homer algorithm (Heinz et al., 2010) to identify enriched *de novo* TF binding motifs within our clustered elements with high statistical stringency (*p*<10-12, Binomial test) (Fig. 5C). We detected trans-regulatory inputs encrypted in *cis*, reflecting chromatin accessibility dynamics, including the binding signatures of early acting factors, canonical NC factors, as well as inputs specifically correlated to either neural or mesenchymal lineages (Fig. 5C). A canonical NC signature highly enriched in *k*-Cluster-3 and late DiffBind elements (Fig. 5C, red box), provided genome-wide evidence of the pivotal role of TFAP2s, *Sox10* and Fox factors in the NC *cis*-regulatory landscapes, suggesting a role for feedforward loops in the global regulation of *bona fide* NC. Moreover, we also found factors suggestive of mesenchymal fates, such as cardiac specifier Pax3, pigment cell driver MITF, myoblast regulator MyoD, and mesenchymal stem regulator NFAT; some of which are known to act downstream of *Sox10*, TFAP2 and FoxD3 (Choi et al., 1990; Conway et al., 1997; Sauka-Spengler and Bronner-Fraser, 2008; Zhu et al., 2014). Amongst other *k*-Cluster-3 motifs, we also found sub-groups of exclusive inputs (e.g., Sox3) and those shared with *k*-Cluster-9 (i.e., ATF, Maf, retinoid acid receptors, Zic, Zfp and GATA) and with *k*-Cluster-6 (Tead1/2 and Meis1/2), possibly reflecting the role for some of these factors in stem cell-like and neural lineages.

The *k*-Cluster-9 motif signature (Fig. 5C, green box) was consistent with the putative role of these elements in neural lineages. For instance, high enrichment in *Sox9* and FoxD3 motifs is in line with the postulated role for these factors in driving the switch from neural to mesenchymal lineages in early trunk NC (Nitzan et al., 2013), but also provides further support for the early heterogeneity of the NC at the cranial level. Intriguingly, we also found significant inputs from Otx2 and Pax2, factors known to play canonical roles in the early neural specification and neural differentiation (Finkelstein and Perrimon, 1991), whose placement within the NC-GRN has been long hypothesised (Le Douarin and Kalcheim, 1999). Moreover, crucial NC nuclear receptors NR2C2 and NR2F2 (Rada-Iglesias et al., 2012) were also found to drive *k*-Cluster-9 elements, as well as an enrichment of SoxC/SoxD factors (Sox4/5), consistent with the role of *k*-Cluster-9 in the early onset of NC programmes. We also uncovered a *k*-Cluster-6 specific niche of upstream inputs including self-renewal genes Oct2/4 and Klf, EMT regulator Zeb1, and the insulator protein CTCF, possibly suggesting that the establishment of NC TADs commences from gastrulation or insulator elements are reused through development. Finally, a signature of premigratory NC associated TFs (e.g., Sox2/9, Msx2, Lef1/Tcf and Pax7) was associated with DiffBind elements at 5-6ss, whereas DiffBind elements at 8-10ss showed the highest enrichment of factors associated with migratory NC (*Sox10*, Sox8, TFAP2s) (Fig. 5C), indicating the role of a Sox8/10-TFAP2-dependent feed-forward loop on the global regulation of migratory NC. Remarkably, these results allow us to conclude that our DiffBind elements indeed capture the temporal dynamics of pre- and migratory *bona fide* NC, whereas *k*-Cluster-3, −6 and −9 elements reveal a regulatory interconnection of canonical and non-canonical NC programmes to different cell-types and lineages, including naive stem cells, neural and mesenchymal progenitors and their lineages. Moreover, enrichment of binding sites for factors not expressed at stages or cell-types studied here (such as anterior Hox genes and nuclear receptors responsive to retinoic acid (RA) signalling), suggest that these enhancers may be involved in the later patterning of the NC through response to external cues, at it commences migration and colonisation of distal sites in the embryo, such as branchial arches (Trainor and Krumlauf, 2000). Such diversity of canonical and novel, lesser characterised TF inputs reflects the broad potential of NC cells and suggests further regulatory refinement is required to direct NC fates.

We next inquired whether this heterogeneous regulatory logic was reflected at the transcriptional level. To this end, we performed single-cell RNA-seq that resulted in 137 deep NC single cell profiles within the narrow dynamic window (6-7ss) that would capture the intrinsic transcriptional dynamics of NC before and during delamination (Fig. 5D). Principle component analysis (PCA) revealed three distinct populations: sc_cluster-1, −2 and −3 (Fig. 5D, E). Sc_cluster-1 and −3 presented clear NC identity, expressing hallmark NC genes (*Pax7, *Sox9*, Snai2, *Sox10*, TFAP2A/B, Ets1*), but segregated according to developmental stage and were uniquely characterised by a number of early-stage specific within sc_cluster-1 (for example, *Brn3, Alx1, Cldn1* and *Sfrp2*,) and later-stage specific, mesenchymal-related NC genes within sc_cluster-3 (for example, *Col9A3, Elk3, Ltk, ItgA9* and *Cxcr4).* In contrast, sc_cluster-2 contained cells collected at both 6ss and 7ss that in addition to sharing some of the NC factors with sc_cluster-1 and −3, also uniquely expressed genes indicative of a neural phenotype, such as *NR2F2, Sox3, CXCL12, Cdh2* etc., including a number of key neural genes exclusively driven only by *k*-cluster-9 elements (*Vimentin, Nbeal1, Pax6* etc) or in concert with *k*-Cluster-6 (*Otx2, Gli2, MylK, Nlgn1*, and retinoic acid signalling players *Nav2, Cyp1b).* Interestingly, while initiating neural differentiation programmes, sc_Cluster-2 cells also expressed a number of NPB specification factors (*Sall1, Zic1, Msx2, Meis1/2, Frzb*, etc.) involved in the maintenance of NC programmes, that were mainly driven by *k*-Cluster-6 elements. Strikingly, sc_Cluster-2 cells did not express majority of canonical, mesenchymal NC markers (Fig. 5D, E and Fig.S7 ShinyApp).

We have demonstrated that the cranial NC cell population at the stages analysed is non-homogenous, albeit not completely resolved to highly divergent clusters that would represent mutually exclusive lineages. However, identified single-cell clusters significantly link to the uncovered heterogeneity at the regulatory level. Premigratory NC sc_Cluster-1 links only to the *k*-Cluster-3 elements (\**p*=0.02), whereas *bona fide* NC sc-cluster-3 is significantly associated to both canonical *k*-Cluster-3 and the early neural *k*-Cluster-9 elements (\*\*\*\**p*=2.23×10^−6^ and \*\**p*=0.003, respectively; two-tailed hypergeometric P-test) (Fig. 5B, E). Strikingly, the singular neural-NC cluster (sc_Cluster-2) was controlled only by early *k*-Cluster-6 and neural-NC *k*-Cluster-9 enhancers (\**p*=0.018 and \*\*\**p*=4.0×10^−4^, respectively) (Fig. 5B, E). Such a strong correlation between *k*-Cluster enhancers and single-cell gene expression clusters indicate the role of early *cis*-regulatory heterogeneity in establishing diverse NC progenitor lineages. Taken together, our results highlight the identification of novel niches of early neural and mesenchymal progenitor NC cells, which are differentially regulated by dynamically accessible elements and their cognate TF inputs with cellular specificity.

### A combinatorial *cis*-regulatory code programmes NC fate restriction

The establishment of regulatory developmental programmes requires combinatorial TF activity (Cunha et al., 2010; Junion et al., 2012; Zinzen et al., 2009). Thus, we next examined putative TF co-binding patterns at NC enhancers, as a means of identifying the core transcriptional networks underlying these distinctive identities. Screening all 2-way combinations of enriched *de novo* TF motifs in our enhancer clusters allowed us to detect a cohort of significantly enriched co-occurring TFs (*p*<0.05, two-sided Chi-squared test with Bonferroni correction) (Fig. 6A, B, Fig. S6).

**Figure 6:**
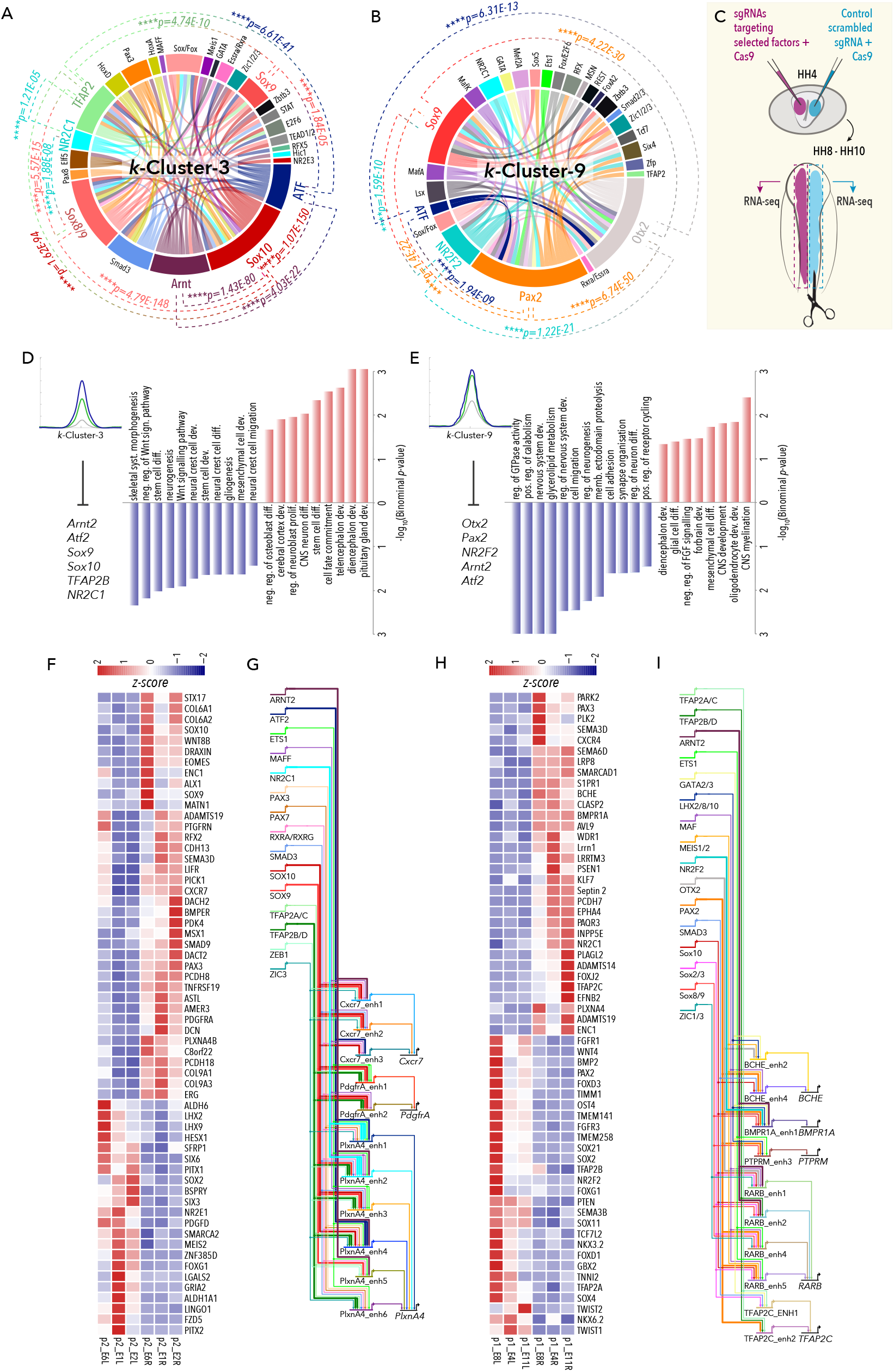
Combinatorial motif analysis and perturbation of the NC-GRN. **(A, B)** TF co-binding relationships predicted in *k*-Cluster-3 (A) and *k*-Cluster-9 (B) elements (B). **(C)** Schematic of bilateral electroporation assay for CRISPR mediated GRN perturbation experiment. **(D, E)** GO terms associated with mis-regulated genes following knock-out of indicated TFs associated with *k*-Cluster-3 (D) and *k*-Cluster-9 (E). **(F)** Heatmap showing relative gene expression changes from left and right dissected dorsal neural tubes from three representative embryos where *k*-Cluster-3 TFs were targeted. (G) Sub-circuitry of gene regulatory interactions governing *Cxcr7, PlxnA4* and *PdgfrA*, all downregulated in *k*-Cluster-3 perturbation experiment. **(H)** Heatmap showing relative gene expression changes from left and right dissected dorsal neural tubes from three representative embryos where *k*-Cluster-9 TFs were targeted. **(I)** Sub-circuitry of gene regulatory interactions governing *BCHE, BMPR1A, PTPRM, RARB* and *TFAP2C*, all downregulated in *k*-Cluster-9 perturbation.

In the *k*-Cluster-3, we found a striking enrichment of canonical interactions specific to *bona fide* NC (Fig. 6A, Fig. S6C), such as *Sox10*-TFAP2 (*p*=1.6×10^−94^), *Sox10*-*Sox9* (*p*=5.8×10^−13^), *Sox10*-Sox8 (*p*=4.8×10^−148^), *Sox10*-Sox/Fox (*p*=7.7×10^−85^) (Sauka-Spengler and Bronner-Fraser, 2008), as well as novel ones, such as ATF-*Sox10* (*p*=1.07 x10^−150^) and Arnt/Arnt2-*Sox10* (*p*=1.4×10^−80^), suggesting a dynamic interplay of TF combinations in the regulation of NC-specific identity. Moreover, analysis of DiffBind at 5ss and 8ss enabled the further identification of temporally restricted, NC-specific combinations controlling premigratory (e.g. Lhx-Sox2, Fox-Sox2, Dmbx1-Lhx, *p* ¡ 2.7×10^−28^) (Figure S6A, C) and early migrating (i.e., *Sox10*-TFAP2s, Smad3-Sox/Fox, *Sox10*-Smad3, TFAP2-Sox/Fox, *p* < 2.3×10^−32^) (Figure S6B, C) identities, respectively (Sauka-Spengler and Bronner-Fraser, 2008). *K*-Cluster-9, however, showed a remarkable enrichment in non-canonical NC TF interactions (Fig. 6B, Figure S6C), such as co-binding of Otx2-Pax2 (*p*=6.7×10^−50^), Otx2-*Sox9* (*p*=4.2×10^−30^), Pax2-*Sox9* (*p*=8.5×10^−24^), NR2F2-Otx2 (*p*=1.4×10^−22^), Otx2-Zic1/2/3 (*p=* 2.9×10^−11^) and NR2F2-*Sox9* (*p*=1.6×10^−10^) amongst others, highlighting and integrating the previously unknown core transcriptional circuitry underlying NC-derived neural progenitors into our current view of the NC-GRN (Betancur et al., 2010a; Sauka-Spengler and Bronner-Fraser, 2008; Simoes-Costa and Bronner, 2015, Sauka-Spengler, 2008). Importantly, these results place Otx2 and Pax2 alongside the canonical NC regulators *Sox9*, Zic1-3, Smad2/3 and NR2F2 (Rada-Iglesias et al., 2012; Sauka-Spengler and Bronner-Fraser, 2008), as novel, core regulators of the NC-GRN, displaying previously unappreciated connectivity to downstream, neural effector genes.

To functionally probe the core transcriptional circuits governing NC heterogeneous identities (canonical/mesenchymal versus neural NC), we used CRISPR-Cas9 (Williams et al., 2018) to endogenously knock out combinations of upstream co-acting factors. Using *in vivo* bilateral co-electroporation assay, we delivered the target gRNAs with wildtype Cas9 to the left and the control gRNA with Cas9 to the right side of the embryo and, following micro-dissection, performed RNA-seq on left and right dorsal neural tubes samples at 7ss, to assess the global effect of the perturbation on the NC-GRN (Fig. 6C, Fig. S7A). On the one hand, we targeted the factors constituting the *k*-Cluster-3 minimal transcriptional core (*Sox10*, *Sox9*, TFAP2B, ATF2, Arnt2 and NR2C1) (Fig. 6A), which we have proposed to underlie NC canonical/mesenchymal progenitor identities (Fig. 5). This strategy caused specific downregulation of 241 and upregulation of 62 genes (FDR<0.1, [LogFoldchange] > 1). Downregulated genes included key premigratory NC regulators such as *Msx1* and *Draxin*, further reinforcing the role of feedback loops in this transcriptional network. Furthermore, downregulation of a number of downstream effector genes (*PlxnA4, Pdgfra, Cxcr7, Col9A1/A3, Pcdh18* and *Cdh13*) (Fig. 6D,F; Fig. S7B), some of which were previously described as functional components of NC programme with roles in cell migration (Olesnicky Killian et al., 2009; Smith and Tallquist, 2010; Waimey et al., 2008), indicated a general disruption of effector gene batteries. Conversely, the upregulated genes were involved in neural programmes (*FoxG1, Hesx1, Lhx2/9, Six3/6, ALDH1A1*) (Aguiar et al., 2014; Andoniadou et al., 2007; Peukert et al., 2011), consistent with the GO terms associated to them (*p*< 0.05, Binomial test with Bonferroni correction) (Fig. 6D).

In a separate experiment, we sought to perturb the factors constituting the *k*-Cluster-9 minimal transcriptional core (Pax2, Otx2, NR2F2, ATF2, and Arnt2) (Fig. 6B) which we proposed to play a critical role within the core transcriptional network driving neural-NC programme (Fig. 3, 5, 6B). GO term analysis of these genes (*p*<0.05, Binomial test with Bonferroni correction) revealed a complex perturbation of a large number of general processes, including catabolism and glycerolipid metabolism, but also downregulation of neural genes (*BCHE, BMPR1a, Lrrn1, PCDH7, Sema6, Cxcr4, PTRM, RARb, TFAP2C, Pax3*) involved in cell migration, neurogenesis, cell adhesion and neural differentiation (Figure 6E, H; Fig. S7C). Interestingly, key TFs and nuclear receptors governing mesenchymal programmes were modestly but significantly upregulated (i.e. *TFAP2A/B, Twist1/2, FoxD1/3, Wnt4, FGFR1/3, Bmp2*), alongside other neural-related pathways dependent on *NKX6.2, Sox2/11/21, PTEN* and *Sema3b*, known to govern neural differentiation (Fig. 6H, Figure S7D). Our results indicate that while perturbing the core transcriptional network of *k*-Cluster-3 causes the direct suppression of NC-specific mesenchymal progenitor programmes with concurrent upregulation of neural signature genes, the disruption of *k*-Cluster-9 regulatory core appears to have an opposite effect on the global transcriptional landscape. Thus, our knockout strategy caused a dysregulation of the NC-specific networks balancing the shift of neural-to-mesenchymal progenitor states, suggesting that the uncovered NC transcriptional cores, mediated by *k*-Cluster-3 and *k*-Cluster-9 regulatory elements balance the shift between NC-derived mesenchymal and neural lineages, likely through mutual cross-negative regulation.

### Integrating regulatory information into GRN circuitry

Current state-of-the-art approaches to reverse engineer global vertebrate GRNs have either used the empirical reconstruction of GRN hierarchies by screening motif binding sites around promoters, together with epistatic information acquired from high-resolution single cell profiles (Aibar et al., 2017), or mathematical modelling of best-fit topologies of epistatic interactions and developmental trajectories determined at the single cell transcriptomic level (Gouti et al., 2017). Here, we build on these strategies by benefiting from the unbiased, high signal-to-noise ratio, low input epigenomes of NC cells and high-resolution single-cell transcriptome information. We unravelled the co-binding dynamics of TFs directly regulating the NC programmes by performing a comprehensive, database-driven screen to identify not only those TFs targeted by our approach, but of all known high-resolution vertebrate motifs (*p*<0.0001, Binomial test) present in our enhancer sets (that is, in the *k*-Clusters −3 and −9, and DiffBind at 5-6ss and 8-10ss) as a means of identifying the full ensemble of TF inputs. We further filtered motif occurrences by integrating TF coexpression data from the single-cell profiles (Fig. 5D, ShinyApp). We next produced comprehensive, genome-wide reconstructions of the *k*-Cluster-3 and *k*-Cluster-9-dependent GRNs (Supp. Files 1 and 2), allowing us to identify the hierarchical position of *k*-Cluster-3 (*Sox10*, *Sox9*, *TFAP2B, ATF2, Arnt2* and *NR2C1*) and *k*-Cluster-9 (*Otx2, Pax2, NR2F2, Atf2* and *Arnt2*) minimal core targeted factors within their respective NC-GRNs, as well to predict their downstream targets.

The extracted GRN circuits that control representative genes downregulated upon *k*-Cluster-3 TF core knockout (Fig. 6D), feature contributions from multiple enhancers within these loci varying between six (*PlxnA4*) to two (*Pdgfra*) per gene (Fig. 6G), further highlighting the importance of enhancer redundancy for the robustness of transcriptional networks. These enhancers primarily mediate heterotypic binding of the core TFs, TFAP2B and *Sox9*/10 as well as additional inputs from other co-binding partners, including Arnt2, NR2C2, MafF, retinoid acid receptors, Zeb1 and Zic3 (Fig. 6G). Some of these inputs form the *k*-Cluster-3 core circuits (Fig. 6A) and are themselves regulated by other targeted factors (NR2C1 and Arnt2, for instance) again highlighting feed-forward loops as core features of the NC-GRN (Supplemental Files 1-5). The extracted GRN circuits controlling representative genes downregulated upon *k*-Cluster-9 TF core knockout (Fig. 6E) identify the direct inputs of disrupted TFs, such as Otx2-Lhx2 and Smad3-Pax2, into BCHE, BMPER1 and TFAP2C, and Otx2-Zic1/3 into RXRb, with additional inputs (NR2F2, Arnt2, Meis1/2, Sox8-10) contributed to individual enhancers (Fig. 6I). These results suggest that the core transcriptional network captured by *k*-Cluster-9 elements acts upstream of RA signalling, through Otx2-dependent activation of RXRb, which may dynamically control the response of these cells to gradients of RA, a critical process in the patterning of neuronal lineages (Gouti et al., 2017). Taken together, our results corroborate the idea that *k*-Cluster-9 elements orchestrate the early establishment of the early neural progenitor state shared between NC and non-NC cells.

Generating sub-circuits governing a small number of genes allows representation of regulatory dynamics and demonstrates that patterns of co-regulation can be resolved for functionally related genes. Moreover, combinatorial patterns of TF inputs suggest the existence of regulatory modules within the broader NC-GRN, possibly related to the co-expressed gene networks that we identified earlier (Figure 1G). As such circuitry requires cellular co-expression of all molecules involved, we used our single-cell profiles to assess co-expression of upstream TF inputs and downstream genes in order to validate our networks further (Fig. S7C). This approach validates novel regulatory relationships and elucidates the core components of the NC-GRN, highlighting the importance of enhancer regulation by combinatorial TF binding for normal NC development.

### Database resource

We assembled our data into a user-friendly, interactive Shiny App (Fig. 7), thus providing a searchable interface whereby individual or multiple genes can be queried for: (1) gene expression; (2) WGCNA cluster association; (3) regulatory elements and upstream regulatory inputs; (4) *k*-cluster dependent transcriptional networks visualisation; and (5) visualisation of single-cell co-expression profiles. This unique tool offers valuable resource for the NC and wider GRN fields, allowing to explore the wealth of novel regulatory sequences and interactions reported here. In-depth analysis of the NC GRN circuits of interest that are easily retrievable using the Shiny App is poised to enable end-to-end investigations of the full complement of NC genes, and thus provide critical insights into precise gene regulatory events underlying NC ontogeny.

**Figure 7:**
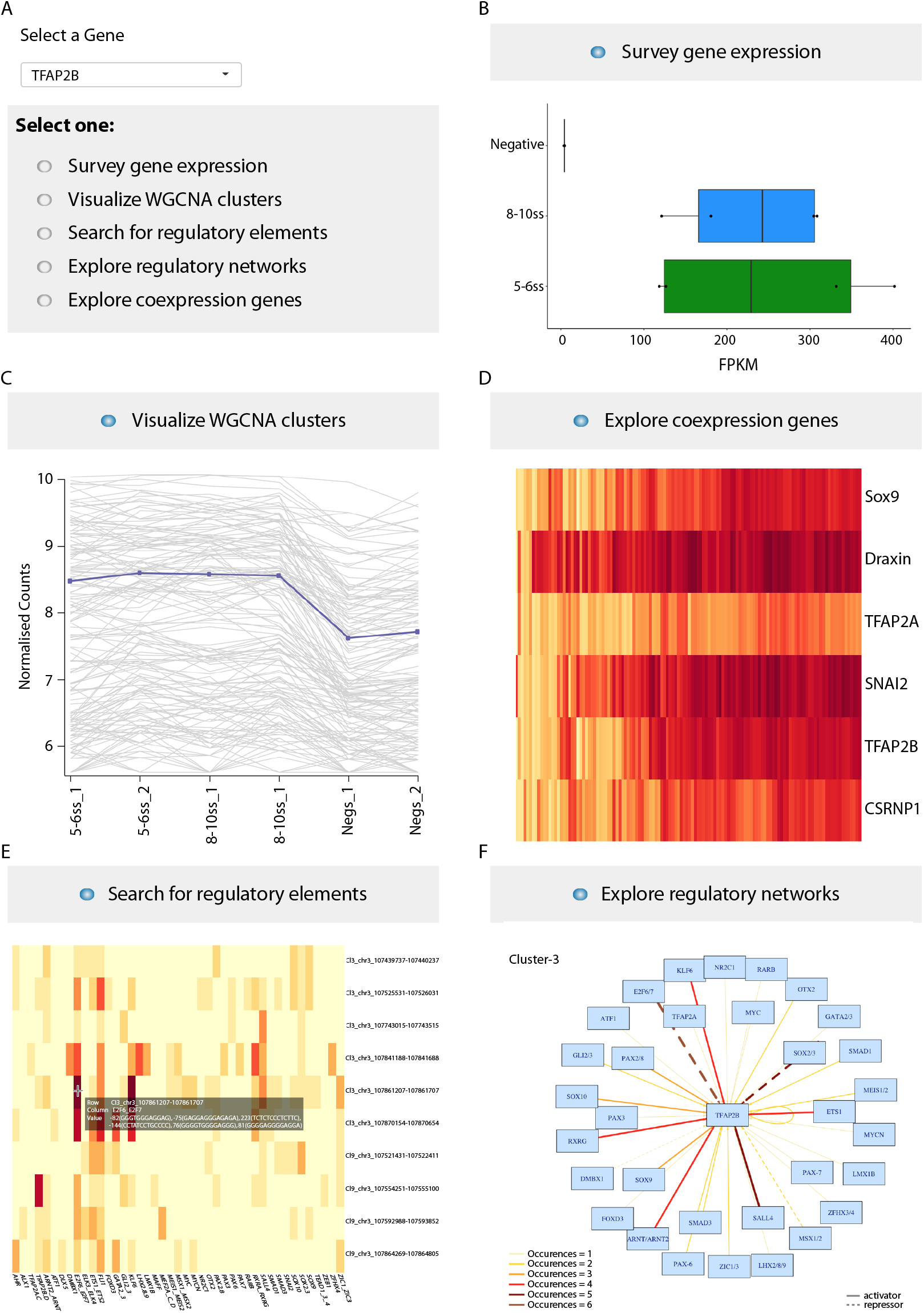
Using Shiny App to explore gene expression and regulation. **(A)** Searchable interface for gene of interest (GOI). **(B)** Gene expression dynamics can be determined across samples analysed, shown in FPKMs, statistically significant enrichment is also shown in the App. **(C)** WGCNA cluster containing GOI can be identified, all other genes present in the same cluster are given and this information is searchable. **(D)** Co-expression with other genes is shown in heatmap form, percentage overlap can be adjusted or groups of specific genes of interest can be selected. **(E)** Associated regulatory elements and inherent TF motifs can be explored. Genome co-ordinates for putative enhancer regions are given for GalGal4 and exact location of motifs within the region is also provided. **(F)** Regulatory networks have been assembled based on *k*-cluster-3 and *k*-Cluster-9 elements and their respective TF inputs. TF occurrence frequency and activating/repressing interactions are shown.

## Discussion

Gene regulatory networks control key regulatory events that ensure accurate development. Over many years of investigation, a wealth of knowledge about NC formation has been accumulated through bottom-up, candidate-gene approach studies, providing essential insights into developmental processes governing NC ontogeny (Le Douarin and Kalcheim, 1999; Sauka-Spengler and Bronner-Fraser, 2008; Bronner and Simoes-Costa, 2016). By investigating the NC gene regulatory mechanisms genome-wide, we provided a top-down, unbiased approach to interrogate global events governing NC ontogeny, allowing us to reverse engineer the comprehensive representation of the NC-GRN. To our knowledge, this represents the first high-resolution, genome-wide reconstruction of a vertebrate GRN *in vivo*, highlighting the potential of our method to decode complex biological networks underlying ontogeny.

### Gene expression signatures of NC

Our transcriptional analysis has yielded the full complement of factors influencing NC formation, both at the cell-autonomous and non-cell autonomous level, significantly increasing the resolution of NC-GRN (Betancur et al., 2010a; Simoes-Costa and Bronner, 2015). For instance, in addition to assigning known and novel regulators to the canonical modules of premigratory (*TFAP2A/B, Msx1, Zeb2*, *Sox9*, *Pax7, Dmbx1* etc.) and migrating NC (*Sox5*, *Sox10*, *Sox8, Lhx9, Pax3, Wnt4, Wnt3a, Ednrb, Robo1, Cxcr4*, etc.), we have also clearly defined a shift from homophilic cell-to-cell communication based on cadherins and protocadherins (*Cdh6/13, Pcdh8*) to heterophilic cell-ECM communication (*Ednrb, Cxcr4, PlxnA4, Adamts1/20*), revealing rewiring of the gene regulatory network as cells undergo EMT and commence migration. Moreover, characterisation of co-expression clusters has expanded the previously proposed modular network structure to refine NC factors into more precise categories (early NC specification, onset of EMT, *bona fide* (late) NC specification, active NC migration, extracellular signalling linked to NC specification, etc.), enabling us to include and classify novel regulators within the NC-GRN. For instance, the zinc-finger homeobox protein 4 (*Zfhx4*), which was recently shown to regulate multipotency, self-renewal and migration of therapy-resistant glioblastomas (NC-derived brain cancer) cells (Chudnovsky et al., 2014), was recovered in the late NC specification module containing core *bona fide* NC specifiers (*Sox10*, *TFAP2A, TFAP2B, Pax7*, etc.). Similarly, another NC regulator - the LIM adaptor protein, *Lmo4* (Ochoa et al., 2012), that featured highly specific NC regulatory control as per our analysis, belonged to the GRN module featuring myocyte-specific enhancer factors *MEF2A/C*, and late non-canonical Wnt factors (*Wnt7A/B*), required in NC for proper craniofacial development and during migration, respectively (De Calisto et al., 2005; Verzi et al., 2007).

### Chromatin dynamics infers NC regulatory architecture

Investigation of NC regulation at the epigenomic level has uncovered dynamic changes in chromatin accessibility of distal regulatory elements employed in the NC and other embryonic cell-types at different stages of NC ontogeny. Our global analysis of chromatin dynamics has first revealed: (1) an NC-independent class of ubiquitously accessible elements, enriched at promoters (*k*-Cluster-4); (2) a class of elements specifically open only in the NC, at either premigratory and/or migrating stages (*k*-Cluster-3); (3) a class of elements dynamically open in both NC and neuroepithelial cells (*k*-Cluster-9); and (4) a class of elements accessible in naive epiblast and premigratory NC cells that become inaccessible by migrating stages (*k*-Cluster-6). Dissection of the specific enhancer clusters by TF binding and co-binding motif enrichment analysis, target gene assignment by positional information and high-resolution targeted chromosome conformation capture analysis of enhancer-promoter contacts, and functional classification of assigned target genes has indicated that three key clusters (*k*-Cluster-3, −6 and −9) embed the regulatory logic of the NC development. This approach has allowed us to not only infer the full complement of active NC enhancers but also to garner the information about the upstream inputs and downstream regulatory outputs from which the NC-GRN could be reconstructed. The *k*-Cluster-3 elements were mainly associated with the *bona fide* NC loci, such as *TFAP2A/B* and *Sox10*, whereas *k*-Cluster-6 and −9 elements were respectively enriched at loci related to stem cells and neural progenitors, such as *Oct4* and *Otx2.* Functional interrogation of multiple *cis*-regulatory elements from the *Sox10* super enhancer region using CRISPR/deadCas9-mediated epigenome editing has revealed the hierarchical contribution of different classes of enhancers to the NC-GRN. As expected, decommissioning individual enhancers caused little effect. Only disruption of the earliest acting element, enh-99, that belonged to the neural-NC *k*-Cluster-9 caused moderate, albeit not sustained disruption of *Sox10* transcription. This suggested that the establishment of the early neural regulatory landscape may be essential for the onset of *bona fide* NC programmes. However, when multiple enhancers were decommissioned, a robust effect was detected, highlighting the role of enhancer redundancy in the robustness of transcriptional networks.

### Elucidating the regulatory basis for NC lineages

By analysing circuitries mediated by *k*-Cluster-3 and *k*-Cluster-9 regulatory elements, we have revealed the heterogeneity of *cis*-regulatory landscapes within the cranial NC cells that was clearly linked to the heterogeneity uncovered at the transcriptional level using single-cell profiling. Our results show that identified single-cell NC clusters featured either canonical neural crest or neural-NC identity, and associated to the corresponding *k*-means clusters with high-statistical support, thus linking their *cis*-regulatory and intrinsic transcriptional dynamics. Our data have thus postulated the existence of an early premigratory NC transcriptional network defining cranial NC fates.

The identified unique combinatorial *cis*-regulatory codes underlying canonical NC identity feature *bona fide* NC interactions *Sox10*-Sox8/9, *Sox10*-TFAP2, as well as co-activities with critical new players ATF and Arnt whereas neural-NC identity is enriched for non-canonical interactions including Otx2-Pax2, Otx2-NR2F2, Pax2-*Sox9* and Otx2-*Sox9*. Our results thus expand a previously proposed minimal network underlying the switch from neural to mesenchymal fates, involving *Sox9* and FoxD3 (Nitzan et al., 2013), showing a previously unappreciated functional role for Otx2, Pax2 and NR2F2 in the NC-GRN. By performing CRISPR/Cas9 knockout of critical top regulators of both programmes *in vivo*, we showed that corresponding transcriptional networks mediating the activation of canonical NC regulators and direct repression of neural pathways balance the neural and mesenchymal NC identities. The activation of an entire *cis*-regulatory landscape specific to canonical NC and mesenchymal progenitors, represented by *k*-Cluster-3 elements, appears vulnerable to perturbation, as the disruption of the corresponding core transcriptional network leads to a robust change towards a neural progenitor state. However, the rewiring of the early neural *cis*-regulatory landscape, which is represented by a large number of strongly interconnected *k*-Cluster-9 elements that are robustly coupled by auto-regulatory and feed-forward loops and active in both NC and neuroepithelial progenitors, appears more difficult to achieve. Such a strong, early relationship between NC and non-NC neuroepithelial programmes may to some extent explain the NC-derived neural progenitors resilience to reprogramming (Simoes-Costa and Bronner, 2016). Nonetheless, the modest yet significant effect in shifting from neural to mesenchymal fates (as revealed by upregulation of *Twist1/2* and *TFAP2a/b*, among others), primarily dictated by *Otx2* and *Pax2* disruption, indeed suggests that these two core networks mutually negatively regulate one another and thus dynamically balance neural and mesenchymal NC lineages.

The involvement of Orthodenticle homeobox 2 (*Otx2*) gene within the NC-GRN has long been hypothesised (Le Douarin and Kalcheim, 1999), given the craniofacial abnormalities that accompanied heterozygous *Otx2*^+/-^ mice (Acampora et al., 1995). Because *Otx2* deficiency causes atrophies and deletions, *Otx2* has thus been proposed to act as a gap gene in vertebrates, in contrast to its function both in homeotic and gap-like transformations in Drosophila (Finkelstein and Perrimon, 1991; Le Douarin and Kalcheim, 1999). Our unbiased, systems-level approach, has enabled the positioning of this factor within the NC-GRN, by identifying its upstream regulators (for example, *Sox9*/10, Sox2/3, Pax2, Msx1/2, Arnt/Arnt2, E2F6, Lhx, RXRg, Pax2/8 and self-regulation) as well as numerous downstream targets in the avian NC crest, (including *Adamts19, Cdh6/9/10/13, Col4a2/6, Ednrb, Epha6/7, Fgfr3, Pcdh8/9/15/19, Pdfrl, PlxnA4, Robo1, Sema3D/5B/6A/6D, Smad3, Sox11, Zfhx4* and *Zic1* among many others). Similarly, the non-canonical role of Pax2 and NR2F2 in controlling neural fate restriction in NC cells illustrates the power of our approach to resolving novel, functionally relevant hierarchies and genome-wide circuits.

### A genome-wide view of the NC-GRN

Our integrated approach using the intermediary of active NC-specific enhancers, has enabled the assembly of comprehensive NC GRNs, linking predicted upstream regulatory factors to their downstream target genes (Supplemental Files 1-5). We have revealed new hierarchies and modular relationships as well as recapitulated previously known interactions, highlighting the utility of our approach to interrogate the complex regulatory logic controlling early NC ontogeny. Moreover, the fact that at least 57% of the genes downregulated upon specific network core perturbations were directly regulated by at least one *k*-Cluster-3 or −9 element as per our analysis confirms the causal links within the proposed NC-GRN circuits. With the recent and on-going development of novel, low input methods to profile TF binding *in vivo* (Skene and Henikoff, 2017), we expect to soon be able to circumvent the key limitation of our approach, which is the reliance on *cis*-regulatory dissection through the use of known, high-resolution motifs.

Our data provide whole-genome framework for understanding the early regulatory events underlying essential NC cell functions. We expect that our resources will be useful for the interrogation of the circuitries underlying essential processes during NC ontogeny, such as maintenance of stemness and multipotency, as well as broader cellular processes and behaviours such as EMT, migration, differentiation and fate restriction.

## Author contributions

Conceptualisation, R.M.W., T.S.S.; Methodology, R.M.W., U.S.; Software, I.C.F., E.R., D.G., J.T.; Validation, R.M.W.; Formal Analysis, I.C.F., R.M.W., T.S.S.; Investigation, R.M.W.; Writing R.M.W., I.C.F., T.S.S.; Writing Review Editing, all authors; Visualisation R.M.W., I.C.F., D.G., E.R., T.S.S.; Supervision, T.S.S.; Funding Acquisition, T.S.S.

## Acknowledgements

We thank all members of the TSS lab for helpful discussions and in particular I. Ling and A. Kenyon for review of the manuscript. FACS was performed at WIMM Flow Cytometry Facility. All next-generation sequencing was performed at the MRC WIMM Sequencing Facility, with the exception of bulk RNA-seq which was sequenced at the Sequencing Core at the WTCHG. High-resolution imaging was conducted in the Wolfson Imaging Centre at the WIMM. This work was supported by MRC (G0902418), The Lister Institute for Preventative Medicine Research Prize, John Fell Fund (131/038), and Leverhulme Trust grant (RPG-2015-026) to T.S.S. I.C.F is funded by the Oxford-Angus McLeod-St John’s College Graduate fellowship and the WIMM prize studentship.

## Data and materials availability

The beta version of the ShinyApp allowing access to the data and network construction can be downloaded from https://github.com/tsslab/chick_NC-GRN. The data and images associated with the ShinyApp can be downloaded from https://figshare.com/articles/chick_NC-GRN_RData/6953294 and https://figshare.com/articles/chick_NC-GRN_images/6953306. The raw and processed data generated in this study have been submitted to GEO (GEO record GSE121527).

## Experimental Procedures

### Embryo culture and electroporations

Fertilised wild-type chicken eggs were obtained from Henry Stewart & Co (Norfolk), staged according to Hamburger and Hamilton (1951) (Hamburger and Hamilton 1951). All experiments were performed on chicken embryos younger than 12 days of development, and as such were not regulated by the Animals (Scientific Procedures) Act 1986. Electroporations were performed as previously described (Sauka-Spengler and Barembaum 2008, Betancur, Bronner-Fraser et al. 2010, Simoes-Costa, McKeown et al. 2012, Williams, Senanayake et al. 2018).

### Cell dissociation and FAC-sorting

Dissected cranial regions from electroporated embryos were dissociated with dispase (1.5mg/ml in DMEM/10mM Hepes pH 7.5) at 37°C for 15min with intermittent pipetting to achieve a single cell suspension and with 0.05% Trypsin at 37°C for a final 3min dissociation step. The reaction was stopped and cells were re-suspended in an excess of Hanks (1XHBSS, 0.25% BSA, 10mM Hepes pH8) buffer. Cells were centrifuged at 500g for 10min, re-suspended in Hanks buffer, passed through a 40-μm cell strainers, and centrifuged at 750g for 10min. Pelleted cells were re-suspended in 500μl Hanks buffer. Fluorescent positive cells were sorted and collected using BD FACS-Aria Fusion. We collected ~300 and ~600 NC cells per embryo at 5-6ss and 8-10ss, respectively.

### Bulk RNA extraction, library preparation and sequencing

FAC-sorted cells were washed with PBS and stored at 80°C in lysis buffer. RNA was extracted using Ambion RNAqueous Micro Total RNA isolation kit (Cat.#AM1931, ThermoFisher Scientific), the integrity was checked using Bioanalyser, and only samples with RIN>7 were used for analysis. Sequencing libraries were prepared using Takara SMARTer low input RNA kit and sequenced using 100bp paired-end reads on the Illumina Hiseq2000 platform. At least three biological replicas for each stage were used for analysis.

### Single cell RNA preparation library preparation and sequencing

Individual NC cells were collected by FACS, cDNA was generated and sequencing libraries were prepared as previously described (Picelli, Faridani et al. 2014). Libraries were sequenced using 50bp single-end reads for 96 cells on the Illumina NextSeq500 platform. A 4×10^7^ dilution of ERCC spike in control was used.

### ATAC, library preparation and sequencing

FAC-sorted cells were lysed (10mM Tris-HCl, pH7.4, 10mM NaCl, 3mM MgCl_2_, 0.1% Igepal) and tagmented using Illumina Nextera DNA kit (FC-121-1030) for 30mins at 37°C. Tagmented DNA was amplified using NEB Next High-Fidelity 2X PCR Master Mix for 11 cycles. Tagmentation efficiency was assessed using Agilent Tapesta-tion. ATAC-seq libraries were sequenced using paired-end 40bp reads on the Illumina NextSeq500 platform. Three biological replicates were obtained for each stage.

### H3K27Ac ChIP, library preparation and sequencing

FAC-sorted cells were cross-linked with 1% formaldehyde for 9mins. Fixation was quenched with 125mM of glycine for 5min. Cross-linker was washed out by 3x pellet washes with 1xPBS ((supplemented with 1X PI (protease inhibitors)), 1mM DTT and 0.2mM PMSF) centrifuging at 2000g for 4mins at 4°CC. Pellets were re-suspended in isotonic nuclei extraction buffer ((NEB: 0.5% NP40, 0.25% Triton-X, 10mM Tris-HCl-pH 7.5, 3mM CaCl_2_, 0.25M sucrose, 1mM DTT, 0.2mM PMSF, 1X PI. Cell nuclei were expulsed with 20 strokes using pestle B in a glass homogeniser, pelleted and washed with 1xPBS (with 1X PI, 1mM DTT and 0.2mM PMSF). Nuclei were lysed in SDS lysis buffer (0.7% SDS, 10mM EDTA, 50mM Tris-HCl (pH 7.5), 1x PI). Cross-linked chromatin was sonicated at 11A 10x (10s on 30s off) followed by 6A 8x (30s on 30s off) using a probe sonicator, to achieve 300-800bp fragments. Pre-blocked Protein A Dynabeads were incubated with antibody (H3K27ac, Abcam Ab4729, Brd4, Bethyl Laboratories A301-985A). Sonicated DNA-protein complexes were applied to beads overnight at 4°C. IgG antibody (Millipore 12-370) was used as control and an input sample was taken. Samples were washed 8x with RIPA buffer (50mM Hepes-KOH (pH 8.0), 500mM LiCl, 1mM EDTA, 1% NP40, 0.7% Na-Deoxycholate, 1x PI) and 1x NaCl TE wash (1x TE, 50mM NaCl) at 4°C. Chromatin was eluted from the beads with SDS ChIP elution buffer (50mM Tris-HCl (pH 7.5), 10mM EDTA, 1%SDS). Cross-linking was reversed overnight at 65°C at 1300 rpm. Cellular RNA was digested with RNaseA (0.2mg/ml) at 37°C for 1 hour, and cellular proteins were removed with Proteinase-K (0.4mg/ml) at 55°C for 1 hour. Samples were purified by standard phenol-chloroform extraction and ethanol precipitation. ChIPd DNA was amplified using published protocol (Adli and Bernstein 2011) for small cell number ChIP, followed by library preparation using NEBNext Ultra DNA^*™*^ library prep kit following manufacturers protocol. ChIP libraries were sequenced using 50bp paired-end reads using Illumina Hiseq2500 platform. Three biological replicates for each stage and antibody were obtained.

### Next Generation Capture-C

Capture-C was performed according to the published protocol (Davies, Telenius et al. 2016), with a few minor adaptions. To collect the material, dorsal neural tubes were dissected from 6-7ss embryos and dissociated in nuclei extraction buffer (NEB, as above), 20 strokes with pestle A in a glass homogeniser. Cells were cross-linked with 2% formaldehyde for 8min. Fixation was quenched with cold 1M glycine. Pelleted cells (1500rpm, 4°C, 5min) were washed with 1xPBS, re-pelleted and re-suspended in lysis buffer (10mM Tris-HCl (pH8), 10mM NaCl, 0.2% Igepal-NP40, 1x cOmplete protease inhibitor (PI)), and left on ice for 20min before pelleting. Supernatant was removed and pellets were stored at −80°C. On the day of the Capture, cross-linked cells were re-suspended in water on ice and homogenised with pestle B, 45 strokes, and centrifuged at 14,000 rpm for 5min. Pelleted cells were re-suspended in water, restriction enzyme buffer and SDS (0.25% final) and incubated at 37°C with 1400 rpm shaking for 1 hour. Triton-X-100 (8.25% final) was added and samples were incubated again with shaking for 1 hour. 3x 125U aliquots of restriction enzyme (DpnII, NEB High-concentration 50,000 U/ml), were added several hours apart, such that samples were digested for 16-24 hours. Restriction enzyme was inactivated by 20min incubation At 65°C. To ligate fragmented DNA, water, ligation buffer and 240 U T4 ligase (high-concentration 30U/μl; Cat.#EL0013, ThermoFisher Scientific) was added and samples were incubated overnight at 16°C with 1400rpm mixing. Samples were uncrosslinked overnight at 65°C in presence of 50μg of Proteinase-K and then treated with 300U RNaseA (Cat.#1119915, Roche) for 30min at 37°C. DNA was extracted using standard Phenol:Chloroform method and ethanol precipitation. Thus obtained 3C libraries were then sonicated to 200bp, using Covaris S220 with the following settings: duty cycle 10%, intensity 5, cycles per burst 200, 6 cycles each of 60s and set mode. Sonicated samples were cleaned using Ampure XP beads, quantified by Qubit (ThermoFisher Scientific) and checked using 2200 Agilent Tapestation System. Libraries were prepared using NEBNext Ultra library prep kit and NEBNext Multiplex oligos for Illumina according to the manufacturers protocol with the exception of using Herculase II fusion enhanced DNA polymerase (Agilent Cat# 600675). Libraries were cleaned up with Ampure XP beads and again assessed using Tapestation and Qubit.

Libraries were prepared for hybridisation by adding 5μg Chicken Hybloc competitor DNA (Applied Genetics Laboratories) to 1.5-2μg 3C library, 1nmol Nimblegen-HE universal oligo, and 1nmol Nimblegen-HE index oligo (specific to the indeces used in library preparation) (Nimblegen SeqCap EZ hybridisation and wash kit, Cat.#05634261001, Roche). The assembled reaction was dried by vacuum centrifuge at 50°C. Hybridisation buffer and component were added and the DNA was carefully reconstituted. Target specific biotinylated oligos were preheated to 47°C in 0.2ml PCR tubes. 3C libraries and blocking oligos were denatured at 95°C for 10min before adding to the preheated biotinylated oligos. The mixture was then incubated at 47°C for 64-72 hours. Hybridised samples were washed and M-270 Streptavidin Dynabeads (Cat. #65305, ThermoFisher Scientific) were prepared according to Nimblegen SeqCap EZ hybridisation and wash kit protocol. Hybridised samples we applied to streptavidin beads and incubated at 47°C for 45min with 600rpm mixing. Washing steps were performed as per kits instructions. Amplification of captured multiplex DNA sample was done using the V2 Nimblegen SeqCap EZ accessory kit (Cat.#07145594001, Roche), followed by an Ampure XP bead clean up. As recommended (Davies, Telenius et al. 2016), to maximise the capture, we repeated the hybridisation reaction (24 hours), as well as subsequent washes and amplification steps. Finally, captured libraries were sequenced using 150bp paired-end reads on the Illumina Miseq platform (300-cycle MiSeq Reagent Kit v2, Cat.#MS-102-2002, Illumina). A full version protocol is available at http://www.tsslab.co.uk/resources.

### Generating Nanotag reporter vectors

pTK Citrine/Cerulean/mCherry reporter vectors (Simoes-Costa, McKeown et al. 2012) were modified to generate an easy-insert integration site enabling highly efficient cloning of putative enhancers using rapid simultaneous ligation/digestion reaction in a single tube. To this end, a *lacZ* expression cassette flanked with restriction sites for the type IIs restriction endonuclease BsmBI (similar to (Cermak, Doyle et al. 2011)) was cloned upstream of theof the minimal Thymidine Kinase (TK) promoter. To enable multiplexing and simultaneous detection of reporter activity from different enhancers, we further modified pTK reporter vectors to include unique barcoded DNA-tags (nanotags) (Nam and Davidson 2012) downstream of fluorescent reporters, within their 3UTR regions and upstream of their polyadenylation signal. We thus generated 48 new Nanotag reporter vectors (16 each for pT*k*Citrine, pT*k*Cerulean and pT*k*mCherry). pTK Nanotag vector collection is available from Addgene (https://www.addgene.org/TatjanLSauka-Spengler/).

### Enhancer cloning and testing

Multiplexed, high-throughput enhancer cloning pipeline is abridged below, and the technical protocol is available at http://www.tsslab.co.uk/resources. Putative enhancers were amplified from chick gDNA using primers containing specific sequence tails (see Table S2) to facilitate subsequent cloning into nanotagged reporter vector using a modified GoldenGate (Engler, Gruetzner et al. 2009) protocol. Briefly, gel-purified amplicons were combined with modified pTK nanotag reporter vector with T4 DNA ligase and BsmBI restriction enzyme and subjected to a cycling reaction that allows simultaneous BsmBI digestion and T4-mediated ligation of amplicon into the reporter vector. Endotoxin-free plasmid preparations (E.Z.N.A. Endo Free Plasmid Mini Kit II, Cat.#D6950-02, Omega Bio-Tek or Qiagen endo-free maxi prep kit, Cat.#12362, Qiagen) were prepared for electroporation. Putative enhancers in reporter plasmids were pooled at 0.2μg/l each (10-12 plasmids including positive, NC1, and negative (short-oligos) controls), and electroporated into the entire epiblast of the early chick gastrula embryos (HH4) (Sauka-Spengler and Barembaum 2008, Simoes-Costa, McKeown et al. 2012, Williams, Senanayake et al. 2018). Embryos were allowed to develop to desired stages using *ex ovo* culture in thin albumin. Cranial regions were dissected and extracted RNA was hybridised with Nanostring reporter code-set oligos designed to detect nanotag transcripts. Nanostring assay was conducted as manufacturers protocol and absolute transcript counts were recorded. Enhancers with a count >50 were electroporated individually at 2μg/μl and fluorescent reporter activity was imaged throughout early NC developmental stages. An extended set of electroporation-ready novel positive NC enhancers cloned into the fluorescent reporters is available from Addgene (https://www.addgene.org/TatjanLSauka-Spengler/).

### Imaging analysis

Embryos were imaged on an Olympus MVX10 stereomicroscope with 2.5X objective using Axio Vision 4.8 software. Zeiss 780 Upright confocal microscope was used for imaging at high cellular resolution

### Cryosectioning and immunostaining

Embryos selected for cryosectioning were fixed in 4% PFA (paraformaldehyde) for 1 hour at room temperature (RT) or at 4°C overnight, then washed 3x 10min in 1x PBS. Embryos were cryoprotected in 15% sucrose/PBS overnight at 4°C, followed by overnight incubation in 7.5% gelatine/15% sucrose /PBS at 37°C. Following ~ 4-hour incubation in 20% gelatine/PBS embryos were embedded in 20% gelatine/PBS, snap frozen in LN2 and stored at −80°C. Cryosectioning was performed at 10μm thickness. Prior to immunocytochemistry, excess gelatine was removed from the slides by a brief rinse in pre-warmed PBS (37°C). Sections were rinsed 3x 5min in PBT (2% DMSO, 0.5% Triton X-100 in PBS), blocked in 10% donkey serum in PBT (block solution) for 1 hour at RT, and incubated overnight at 4°C with primary antibody (1:200 dilution of rabbit anti-GFP in block solution, Cat.#TP401, Torrey Pines Biolabs). Sections were then washed in PBT at RT 5x 10min, followed by incubation with secondary antibody (AlexaFluor-488-conjugated donkey anti-rabbit IgG diluted at 1:1000 in PBT, Cat.#A21206, ThermoFisher Scientific) for 2 hours at RT. Sections were washed 6-8x 10min in PBT at RT then overnight at 4°C and mounted using Vectashield with DAPI (Cat.#H-1200, Vector Laboratories).

### Hybridisation chain reaction fluorescent *in situ*

*In situ* v3 HCR for *Sox10* was performed as previously described (Choi, Schwarzkopf et al. 2018, Williams, Senanayake et al. 2018).

### *In vivo* CRISPR-mediated perturbation assays

sgRNAs were cloned as previously described (Williams, Senanayake et al. 2018) and used in conjunction with dCas9-Krab (Addgene #92361) (Williams, Senanayake et al. 2018) to decommission endogenous enhancers or Cas9-2A-Citrine (Addgene #92358) (Williams, Senanayake et al. 2018) to knockout transcription factors. Following bilateral electroporation, left and right dorsal neural tubes were dissected separately; RNA was extracted using Ambion RNAqueous Micro Total RNA isolation kit (Cat.#AM1931, ThermoFisher Scientific). For enhancer decommissioning experiments, oligo-dT-primed cDNA was synthesised using Superscript III (Cat.# 18080093, ThermoFisher Scientific) and qPCR performed using Fast SYBR Green reagent (Cat.# 4385612, ThermoFisher Scientific) on the Applied Biosystems 7500 Fast Real-Time PCR System. The standard curve method was used to quantify gene expression, ~10 embryos per targeted enhancer were analysed. For TF knockout experiments RNA-seq libraries were prepared using SMART-Seq^™^ v4 Ultra^™^ Low Input RNA Kit for Sequencing (Cat.#634889, Takara Bio Clontech) and sequenced using 40bp paired-end reads on Illumina NextSeq500. 6 individual embryos and associated RNA-sew libraries were analysed.

## Statistical Analysis and Bioinformatics data processing

Sequencing files from each sequencing lane were de-multiplexed and the resulting files were merged. Reads were trimmed for quality using sickle (Joshi 2011) (v.1.33) quality control was performed using fastQC (v.0.10.1) and deeptools (Ramirez, Ryan et al. 2016) (v.2.2.2). Reads were mapped to the chicken genome galGal4 assembly using bowtie (Langmead and Salzberg 2012) (v.1.0.0), except for RNA-seq files, which were mapped using RNA-STAR (2.4.2a) (Dobin, Davis et al. 2013). In the CRISPR knock out experiments, reads were mapped to galGal5. Sequence alignment/map files were compressed to the binary version (BAM) for downstream analysis. Differential expression analysis was carried out using in DESeq2 (Love, Huber et al. 2014) (v.1.14.1). Peaks were called for each ATAC-seq sample using MACS2 (v.2.0.10) (Feng, Liu et al. 2012) and ATAC-seq consensus peaks were generated using bedtools (Langmead, Trapnell et al. 2009) (v. 2.25.0) intersect command. For RNA- and ATAC-seq reproducibility analyses, BAM files were first normalized for sequencing depth using samtools (Li, Handsaker et al. 2009) (v.1.3) and count tables were retrieved using Subread (Liao, Smyth et al. 2013) (v.1.4.5) featureCounts (Liao, Smyth et al. 2014) tool from an Ensembl gene list in GTF format (for RNA-seq) and from a consensus set of 99,583 accessible regions (for ATAC-seq), respectively. Count tables were analysed in R (v.3.2.3) and principal component analysis (PCA) was performed using the DEseq2 (v1.4.5) package in the R (v.3.2.1) environment. A custom Perl script was used to generate smoothened genome browser tracks in BigWig format for ATAC-seq data visualisation.

### WGCNA analysis

Weighted correlation network analysis (WGCNA) (Langfelder and Horvath 2008) was performed on normalised gene count tables generated by DESeq2, according to the pipeline detailed in the online tutorial (https://labs.genetics.ucla.edu/horvath/CoexpressionNetwork/Rpackages/WGCNA/).

### Single-Cell RNA-Seq Analysis

Reads were mapped using RNA-STAR (2.4.2a) (Dobin, Davis et al. 2013) to the chicken genome Galgal4 assembly and the ERCC spike-in controls using the default parameters. Gene expression levels were quantified as read counts using the featureCounts function from the Subread package with default parameters. Reads that aligned to more than one locus as well as ambiguous fragments were excluded from all further analysis. To remove cells with low quality sequencing, cells with a) less than 100,000 sequenced reads or b) less than 50% uniquely mapped reads were excluded from any further analysis. Further filtering was done based on the distributions of a) the number of expressed genes per cell, b) the proportion of reads on mitochondrial genes or c) the proportion of reads on ERCC spike-ins, requiring that the cells are within 3 MADs (Median Absolute Deviation) for each distribution (McCarthy, Campbell et al. 2017). In the remaining 124 cells, 10,291 genes were detected using the following criteria: a) they had any reads in more than 5 cells and b) had an average number of counts above 1. The cell-based factors from the deconvolved pool-based size factors were used for the normalisation of the gene counts, as described in (Lun, Bach et al. 2016), using pool sizes of 10, 20,40 and 60 cells. The clustering was done with the pagoda2 package (https://github.com/hms-dbmi/pagoda2) using the infomap and the walktrap community methods. There was a 96% concordance between the two methods and the infomap method was used to obtain the final clusters. An analysis for differentially expressed genes was performed using the SCDE package (Kharchenko, Silberstein et al. 2014) to find cluster specific gene markers. For each cluster a one-versus-one approach was applied between all pairs of clusters using the default parameters.

### K-means clustering

Clustering of downsampled ATAC-seq datasets was performed with the seqMINER platform (v.1.3.4) (Ye, Krebs et al. 2011) on a consensus set of 99,583 accessible regions using the *k*-means enrichment linear clustering normalisation algorithm set to k=10 and a window of ±1.0 kb from the peak centre. Heatmaps and merged profiles were plotted using the Deeptools (v.2.2.2) package (Ramirez, Ryan et al. 2016). *K*-Cluster-specific count tables were then used for correlation and linear regression analyses and visualization in R (v.3.2.3).

### Differential accessibility analysis

Differential accessibility analysis was carried out in R (v.3.2.1) using the DiffBind package (v1.10.2). Differential accessibility across samples was calculated using a negative binomial distribution model implemented in DEseq2 (v1.4.5). A stringent threshold (FDR<0.1, Fold enrichment >1) was used to define high-confidence differentially accessible ATAC-seq peaks by comparing the enrichment of accessible regions in NC cells at two stages (5-6ss and 8-10ss) over non-NC cells. Normalised read counts were clustered and visualised using pheatmap (v. 1.0.10) (Kolde 2018) with default settings.

### Next-Generation Capture-C analysis

NG Capture-C was performed as described previously (Davies, Telenius et al. 2016) on dissected dorsal neural tube tissue and chicken erythrocytes. Samples were indexed for multiplexing and co-capture of promoters using biotinylated 120-mers (Integrated DNA technologies, IDT), designed with the CapSequm webtool (http://apps.molbiol.ox.ac.uk/CaptureC/cgi-bin/CapSequm.cgi) (Hughes, Roberts et al. 2014) and pooled to a final concentration of 2.9nM (Table S1). Captured material was pooled and sequenced using 150bp paired-end reads on the Illumina Miseq platform (300-cycle MiSeq Reagent Kit v2, Cat.#MS-102-2002, Illumina). Reads were mapped using Capture-C scripts (https://github.com/Hughes-Genome-Group/CCseqBasicF/releases), and analysed as previously described (Hay, Hughes et al. 2016) using intra-house R-scripts and DESeq2.

### Functional annotation of CREs

A custom Python (v.2.7.5) script was used to assign peaks to nearest expressed genes. Ensembl identifiers were converted to gene names using the Chicken Ensembl Gene ID converter (https://www.biotools.fr/). Genes associated to two or more enhancers were used for functional analysis and GRN assembly. High-resolution TADs were determined as described in (Davies, Telenius et al. 2016). The Panther package (Mi, Huang et al. 2017) (v.11) was used to calculate statistical overrepresentation tests using default settings. P-values were calculated with binomial distributions with Bonferroni correction for multiple hypothesis testing.

### Motif analysis

*De novo* motif discovery was performed using Homer (Heinz, Benner et al. 2010) (v.4.7) findMotifsGenome.pl script, using clusters of ATAC-seq peaks identified by differential accessibility analysis and *k*-means clustering. Motifs of 822bp in length were searched in ±250bp windows from the peak centre and a constant background containing a set of 99,583 peaks was used. Non-redundant matrices with *p*< 11011 (Binomial P-test) were retained. *De novo* motifs were annotated using Homer (v.4.7), STAMP (Mahony and Benos 2007), TOMTOM (Gupta, Stamatoyannopoulos et al. 2007) and manual comparisons. Log10 transformed P-values were clustered and visualised using pheatmap (v. 1.0.10) in R (v. 3.2.1) with default settings. subsection*Combinatorial binding All possible pairs of cluster-specific *de novo* motif combinations were computed in R (v. 3.2.1). Positive controls were defined by using a set of 99,583 consensus peaks subtracted to each cluster of elements using bedtools (v. 2.25.0) intersect v command. Homer (v.4.7) annotatePeaks.pl script was used to screen all *de novo* motifs occurring in windows of ±250bp from the peak centre. Motif occurrences were converted to a matrix of motif presence (1) or absence (0) in R (v.3.2.3), and a custom Python3 script using the Pandas package was used to calculate motif co-occurrences. Combinations enriched at α= 5% (twotailed Chi-squared test) with Bonferroni correction for multiple hypothesis (m) testing were retained for *P*-values < α/m (Noble 2009). Motif combination frequencies in differentially accessible and in *k*-Clusters groups of elements versus that of positive controls were used as a metric of combinatorial enrichment. Co-binding networks were plotted using the Circlize package in R (v.3.2.3).

### Identification and ranking of super-enhancer-like clusters

Super-enhancers were identified using the ROSE (Ranked Order of Super Enhancers) pipeline (http://younglab.wi.mit.edu/super_enhancer_code.html). This algorithm groups (stitches) identified individual enhancers positioned within a defined distance into putative super-enhancers (SEs) and ranks them by their input-subtracted H3K27Acetylation signal. We used the default options with 12.5 kb as the maximum grouping (stitching) distance, the reads per million normalization for H3K27Ac ChIP signal from 5-6ss and 8-10ss FAC-sorted neural crest and a promoter exclusion zone of 1,000 or 2,500 basepairs. The enhancer clusters were ranked by cumulative normalised H3K27Ac signal (H3K27ac signal versus enhancer rank) and super-enhancers were identified as clusters lying beyond the identified inflection point on the ranking curve (Whyte, Orlando et al. 2013). Super-enhancers cluster were associated to the NC enhancers identified in this study and assigned to the closest expressed gene(s). The identification of groups of NC enhancers enriched in Brd4 occupancy on was achieved by *k*-means clustering (Seqminer platform) (Ye, Krebs et al. 2011) of the Brd4 ChIP datasets using 8-10ss DiffBind enhancers as a reference.

### GRN assembly

A comprehensive database collection of high-resolution TF binding models from vertebrates (HOCOMOCO v.11) (Kulakovskiy, Vorontsov et al. 2018) was downloaded in Homer (v.4.7) format. Position weight matrixes (PWMs) with a stringent Binomial 3-value <0.0001 were selected, resulting in a list of 769 PWMs. These TF binding models were first filtered based on PWM redundancy and were further eliminated by applying a minimum expression level threshold (1>FPKM) to their cognate TFs, resulting in a list of 169 motifs. scRNA-seq was used to define TFs co-expressed in clusters of single cells. This list was further refined by eliminating ubiquitous TFs and by combining TF paralogs (e.g. TFAP2A and C; TFAP2B and D etc. based on high similarity of their binding motifs). This resulted in a final list of 49 PWMs being retained. Homer (v.4.7) annotatePeaks.pl script was used to screen these PWMs in all differentially accessible and in *k*-Clusters groups of elements, allowing the inference of the upstream (TF) inputs putatively binding to such elements. Matrices containing the list of CREs and upstream TF inputs were merged with another matrix containing the CREs assignments to their target promoters, which were determined either by Capture-C or by positional information from expressed genes. These elements were grouped based in the differential accessibility analysis and *k*-means clustering, which facilitated the retaining of the GRN spatiotemporal dynamics. The combination of such matrixes linked the upstream (TF) inputs and the downstream outputs (target genes), resolving GRN hierarchies and modules in a genome-wide fashion, at single-cell resolution and with epigenomic details depicted. GRN hierarchical relationships were assembled and used for network analysis and visualization in BioTapestry (Longabaugh, Davidson et al. 2005) (v. 7.1.1). Sub-networks for specific genes or groups of genes were retrieved from the matrixes containing all hierarchies and can be interactively surveyed in the ShinyApp.

### Data availability

The beta version of the ShinyApp allowing access to the data and network construction can be downloaded from https://github.com/tsslab/chick_NC-GRN. The data and images associated with the ShinyApp can be downloaded from https://figshare.com/articles/chick_NC-GRN_RData/6953294 and https://figshare.com/articles/chick_NC-GRN_images/6953306. The raw and processed data generated in this study have also been submitted to GEO (GEO record GSE121527).

## References

Acampora, D., Mazan, S., Lallemand, Y., Avantaggiato, V., Maury, M., Simeone, A., and Brulet, P. (1995). Forebrain and midbrain regions are deleted in Otx2-/-mutants due to a defective anterior neuroectoderm specification during gastrulation. Development 121, 3279–3290.

Adli, M., and Bernstein, B.E. (2011). Whole-genome chromatin profiling from limited numbers of cells using nano-ChIP-seq. Nat Protoc 6, 1656–1668.

Aguiar, D.P., Sghari, S., and Creuzet, S. (2014). The facial neural crest controls fore- and midbrain patterning by regulating Foxg1 expression through Smad1 activity. Development 141, 2494–2505.

Aibar, S., Gonzalez-Blas, C.B., Moerman, T., Huynh-Thu, V.A., Imrichova, H., Hulselmans, G., Rambow, F., Marine, J.C., Geurts, P., Aerts, J., et al. (2017). SCENIC: single-cell regulatory network inference and clustering. Nat Methods 14, 1083–1086.

Andoniadou, C.L., Signore, M., Sajedi, E., Gaston-Massuet, C., Kelberman, D., Burns, A.J., Itasaki, N., Dattani, M., and Martinez-Barbera, J.P. (2007). Lack of the murine homeobox gene Hesx1 leads to a posterior transformation of the anterior forebrain. Development 134, 1499–1508.

Baggiolini, A., Varum, S., Mateos, J.M., Bettosini, D., John, N., Bonalli, M., Ziegler, U., Dimou, L., Clevers, H., Furrer, R., et al. (2015). Premigratory and migratory neural crest cells are multipotent in vivo. Cell Stem Cell 16, 314–322.

Barembaum, M., and Bronner, M.E. (2013). Identification and dissection of a key enhancer mediating cranial neural crest specific expression of transcription factor, Ets-1. Dev Biol 382, 567–575.

Basch, M.L., Bronner-Fraser, M., and Garcia-Castro, M.I. (2006). Specification of the neural crest occurs during gastrulation and requires Pax7. Nature 441, 218–222.

Betancur, P., Bronner-Fraser, M., and Sauka-Spengler, T. (2010a). Assembling neural crest regulatory circuits into a gene regulatory network. Annu Rev Cell Dev Biol 26, 581–603.

Betancur, P., Bronner-Fraser, M., and Sauka-Spengler, T. (2010b). Genomic code for Sox10 activation reveals a key regulatory enhancer for cranial neural crest. Proc Natl Acad Sci U S A 107, 3570–3575.

Boeva, V., Louis-Brennetot, C., Peltier, A., Durand, S., Pierre-Eugene, C., Raynal, V., Etchevers, H.C., Thomas, S., Lermine, A., Daudigeos-Dubus, E., et al. (2017). Heterogeneity of neuroblastoma cell identity defined by transcriptional circuitries. Nat Genet 49, 1408–1413.

Bothma, J.P., Garcia, H.G., Ng, S., Perry, M.W., Gregor, T., and Levine, M. (2015). Enhancer additivity and non-additivity are determined by enhancer strength in the Drosophila embryo. Elife 4.

Bronner, M.E., and Simoes-Costa, M. (2016). The Neural Crest Migrating into the Twenty-First Century. Curr Top Dev Biol 116, 115–134.

Bronner-Fraser, M., and Fraser, S.E. (1988). Cell lineage analysis reveals multipotency of some avian neural crest cells. Nature 335, 161–164.

Buenrostro, J.D., Giresi, P.G., Zaba, L.C., Chang, H.Y., and Greenleaf, W.J. (2013). Transposition of native chromatin for fast and sensitive epigenomic profiling of open chromatin, DNA-binding proteins and nucleosome position. Nat Methods 10, 1213–1218.

Buitrago-Delgado, E., Nordin, K., Rao, A., Geary, L., and LaBonne, C. (2015). NEURODEVELOPMENT. Shared regulatory programs suggest retention of blastula-stage potential in neural crest cells. Science 348, 1332–1335.

Cannavo, E., Khoueiry, P., Garfield, D.A., Geeleher, P., Zichner, T., Gustafson, E.H., Ciglar, L., Korbel, J.O., and Furlong, E.E. (2016). Shadow Enhancers Are Pervasive Features of Developmental Regulatory Networks. Curr Biol 26, 38–51.

Carter, T.C., Kay, D.M., Browne, M.L., Liu, A., Romitti, P.A., Kuehn, D., Conley, M.R., Caggana, M., Druschel, C.M., Brody, L.C., et al. (2012). Hirschsprung’s disease and variants in genes that regulate enteric neural crest cell proliferation, migration and differentiation. J Hum Genet 57, 485–493.

Cermak, T., Doyle, E.L., Christian, M., Wang, L., Zhang, Y., Schmidt, C., Baller, J.A., Somia, N.V., Bogdanove, A.J., and Voytas, D.F. (2011). Efficient design and assembly of custom TALEN and other TAL effector-based constructs for DNA targeting. Nucleic Acids Res 39, e82.

Choi, H.M.T., Schwarzkopf, M., Fornace, M.E., Acharya, A., Artavanis, G., Stegmaier, J., Cunha, A., and Pierce, N.A. (2018). Third-generation in situ hybridization chain reaction: multiplexed, quantitative, sensitive, versatile, robust. Development 145.

Choi, J., Costa, M.L., Mermelstein, C.S., Chagas, C., Holtzer, S., and Holtzer, H. (1990). MyoD converts primary dermal fibroblasts, chondroblasts, smooth muscle, and retinal pigmented epithelial cells into striated mononucleated myoblasts and multinucleated myotubes. Proc Natl Acad Sci U S A 87, 7988–7992.

Chudnovsky, Y., Kim, D., Zheng, S., Whyte, W.A., Bansal, M., Bray, M.A., Gopal, S., Theisen, M.A., Bilodeau, S., Thiru, P., et al. (2014). ZFHX4 interacts with the NuRD core member CHD4 and regulates the glioblastoma tumor-initiating cell state. Cell Rep 6, 313–324.

Conway, S.J., Henderson, D.J., and Copp, A.J. (1997). Pax3 is required for cardiac neural crest migration in the mouse: evidence from the splotch (Sp2H) mutant. Development 124, 505–514.

Cunha, P.M., Sandmann, T., Gustafson, E.H., Ciglar, L., Eichenlaub, M.P., and Furlong, E.E. (2010). Combinatorial binding leads to diverse regulatory responses: Lmd is a tissue-specific modulator of Mef2 activity. PLoS Genet 6, e1001014.

Davidson, E.H. (2010). Emerging properties of animal gene regulatory networks. Nature 468, 911–920.

Davidson, E.H., and Erwin, D.H. (2006). Gene regulatory networks and the evolution of animal body plans. Science 311, 796–800.

Davies, J.O., Telenius, J.M., McGowan, S.J., Roberts, N.A., Taylor, S., Higgs, D.R., and Hughes, J.R. (2016). Multiplexed analysis of chromosome conformation at vastly improved sensitivity. Nat Methods 13, 74–80.

De Calisto, J., Araya, C., Marchant, L., Riaz, C.F., and Mayor, R. (2005). Essential role of non-canonical Wnt signalling in neural crest migration. Development 132, 2587–2597.

de Laat, W., and Duboule, D. (2013). Topology of mammalian developmental enhancers and their regulatory landscapes. Nature 502, 499–506.

Dobin, A., Davis, C.A., Schlesinger, F., Drenkow, J., Zaleski, C., Jha, S., Batut, P., Chaisson, M., and Gingeras, T.R. (2013). STAR: ultrafast universal RNA-seq aligner. Bioinformatics 29, 15–21.

Dupin, E., Calloni, G.W., Coelho-Aguiar, J.M., and Le Douarin, N.M. (2018). The issue of the multipotency of the neural crest cells. Dev Biol.

Engler, C., Gruetzner, R., Kandzia, R., and Marillonnet, S. (2009). Golden gate shuffling: a one-pot DNA shuffling method based on type IIs restriction enzymes. PLoS One 4, e5553.

Feng, J., Liu, T., Qin, B., Zhang, Y., and Liu, X.S. (2012). Identifying ChIP-seq enrichment using MACS. Nat Protoc 7, 1728–1740.

Finkelstein, R., and Perrimon, N. (1991). The molecular genetics of head development in Drosophila melanogaster. Development 112, 899–912.

Georgescu, C., Longabaugh, W.J., Scripture-Adams, D.D., David-Fung, E.S., Yui, M.A., Zarnegar, M.A., Bolouri, H., and Rothenberg, E.V. (2008). A gene regulatory network armature for T lymphocyte specification. Proc Natl Acad Sci U S A 105, 20100–20105.

Gonen, N., Futtner, C.R., Wood, S., Garcia-Moreno, S.A., Salamone, I.M., Samson, S.C., Sekido, R., Poulat, F., Maatouk, D.M., and Lovell-Badge, R. (2018). Sex reversal following deletion of a single distal enhancer of Sox9. Science 360, 1469–1473.

Gouti, M., Delile, J., Stamataki, D., Wymeersch, F.J., Huang, Y., Kleinjung, J., Wilson, V., and Briscoe, J. (2017). A Gene Regulatory Network Balances Neural and Mesoderm Specification during Vertebrate Trunk Development. Dev Cell 41, 243–261 e247.

Gupta, S., Stamatoyannopoulos, J.A., Bailey, T.L., and Noble, W.S. (2007). Quantifying similarity between motifs. Genome Biol 8, R24.

Hamburger, V., and Hamilton, H.L. (1951). A series of normal stages in the development of the chick embryo. J Morphol 88, 49–92.

Hay, D., Hughes, J.R., Babbs, C., Davies, J.O.J., Graham, B.J., Hanssen, L., Kassouf, M.T., Marieke Oudelaar, A.M., Sharpe, J.A., Suciu, M.C., et al. (2016). Genetic dissection of the alpha-globin super-enhancer in vivo. Nat Genet 48, 895–903.

Heinz, S., Benner, C., Spann, N., Bertolino, E., Lin, Y.C., Laslo, P., Cheng, J.X., Murre, C., Singh, H., and Glass, C.K. (2010). Simple combinations of lineage-determining transcription factors prime cis-regulatory elements required for macrophage and B cell identities. Mol Cell 38, 576–589.

Hnisz, D., Abraham, B.J., Lee, T.I., Lau, A., Saint-Andre, V., Sigova, A.A., Hoke, H.A., and Young, R.A. (2013). Super-enhancers in the control of cell identity and disease. Cell 155, 934–947.

Hughes, J.R., Roberts, N., McGowan, S., Hay, D., Giannoulatou, E., Lynch, M., De Gobbi, M., Taylor, S., Gibbons, R., and Higgs, D.R. (2014). Analysis of hundreds of cis-regulatory landscapes at high resolution in a single, high-throughput experiment. Nat Genet 46, 205–212.

Joshi, N.a.F., J. (2011). Sickle: A sliding-window, adaptive, quality-based trimming tool for FastQ files. available at https://githubcom/najoshi/sickle.

Junion, G., Spivakov, M., Girardot, C., Braun, M., Gustafson, E.H., Birney, E., and Furlong, E.E. (2012). A transcription factor collective defines cardiac cell fate and reflects lineage history. Cell 148, 473–486.

Karunasena, E., McIver, L.J., Bavarva, J.H., Wu, X., Zhu, H., and Garner, H.R. (2015). ‘Cut from the same cloth’: Shared microsatellite variants among cancers link to ectodermal tissues-neural tube and crest cells. Oncotarget 6, 22038–22047.

Kelsh, R.N. (2006). Sorting out Sox10 functions in neural crest development. Bioessays 28, 788–798.

Kharchenko, P.V., Silberstein, L., and Scadden, D.T. (2014). Bayesian approach to single-cell differential expression analysis. Nat Methods 11, 740–742.

Kolde, R. (2018). pheatmap: Pretty Heatmaps. R package version 1.0.10.

Kulakovskiy, I.V., Vorontsov, I.E., Yevshin, I.S., Sharipov, R.N., Fedorova, A.D., Rumynskiy, E.I., Medvedeva, Y.A., Magana-Mora, A., Bajic, V.B., Papatsenko, D.A., et al. (2018). HOCOMOCO: towards a complete collection of transcription factor binding models for human and mouse via large-scale ChIP-Seq analysis. Nucleic Acids Res 46, D252–D259.

Langfelder, P., and Horvath, S. (2008). WGCNA: an R package for weighted correlation network analysis. BMC Bioinformatics 9, 559.

Langfelder, P., Zhang, B., and Horvath, S. (2008). Defining clusters from a hierarchical cluster tree: the Dynamic Tree Cut package for R. Bioinformatics 24, 719–720.

Langmead, B., and Salzberg, S.L. (2012). Fast gapped-read alignment with Bowtie 2. Nat Methods 9, 357–359.

Langmead, B., Trapnell, C., Pop, M., and Salzberg, S.L. (2009). Ultrafast and memory-efficient alignment of short DNA sequences to the human genome. Genome Biol 10, R25.

Le Douarin, N.M., and Kalcheim, C. (1999). The Neural Crest. 2nd ed (New York: Cambridge University Press).

Le Lievre, C.S., Schweizer, G.G., Ziller, C.M., and Le Douarin, N.M. (1980). Restrictions of developmental capabilities in neural crest cell derivatives as tested by in vivo transplantation experiments. Dev Biol 77, 362–378.

Lee, T.I., Rinaldi, N.J., Robert, F., Odom, D.T., Bar-Joseph, Z., Gerber, G.K., Hannett, N.M., Harbison, C.T., Thompson, C.M., Simon, I., et al. (2002). Transcriptional regulatory networks in Saccharomyces cerevisiae. Science 298, 799–804.

Levine, M., Cattoglio, C., and Tjian, R. (2014). Looping back to leap forward: transcription enters a new era. Cell 157, 13–25.

Levine, M., and Davidson, E.H. (2005). Gene regulatory networks for development. Proc Natl Acad Sci U S A 102, 4936–4942.

Li, H., Handsaker, B., Wysoker, A., Fennell, T., Ruan, J., Homer, N., Marth, G., Abecasis, G., Durbin, R., and Genome Project Data Processing, S. (2009). The Sequence Alignment/Map format and SAMtools. Bioinformatics 25, 2078–2079.

Liao, Y., Smyth, G.K., and Shi, W. (2013). The Subread aligner: fast, accurate and scalable read mapping by seed-and-vote. Nucleic Acids Res 41, e108.

Liao, Y., Smyth, G.K., and Shi, W. (2014). featureCounts: an efficient general purpose program for assigning sequence reads to genomic features. Bioinformatics 30, 923–930.

Longabaugh, W.J., Davidson, E.H., and Bolouri, H. (2005). Computational representation of developmental genetic regulatory networks. Dev Biol 283, 1–16.

Love, M.I., Huber, W., and Anders, S. (2014). Moderated estimation of fold change and dispersion for RNA-seq data with DESeq2. Genome Biol 15, 550.

Loven, J., Hoke, H.A., Lin, C.Y., Lau, A., Orlando, D.A., Vakoc, C.R., Bradner, J.E., Lee, T.I., and Young, R.A. (2013). Selective inhibition of tumor oncogenes by disruption of super-enhancers. Cell 153, 320–334.

Lukoseviciute, M., Gavriouchkina, D., Williams, R.M., Hochgreb-Hagele, T., Senanayake, U., Chong-Morrison, V., Thongjuea, S., Repapi, E., Mead, A., and Sauka-Spengler, T. (2018). From Pioneer to Repressor: Bimodal foxd3 Activity Dynamically Remodels Neural Crest Regulatory Landscape In Vivo. Dev Cell 47, 608–628 e606.

Lun, A.T., Bach, K., and Marioni, J.C. (2016). Pooling across cells to normalize single-cell RNA sequencing data with many zero counts. Genome Biol 17, 75.

Mahony, S., and Benos, P.V. (2007). STAMP: a web tool for exploring DNA-binding motif similarities. Nucleic Acids Res 35, W253–258.

Mayran, A., Khetchoumian, K., Hariri, F., Pastinen, T., Gauthier, Y., Balsalobre, A., and Drouin, J. (2018). Pioneer factor Pax7 deploys a stable enhancer repertoire for specification of cell fate. Nat Genet 50, 259–269.

McCarthy, D.J., Campbell, K.R., Lun, A.T., and Wills, Q.F. (2017). Scater: pre-processing, quality control, normalization and visualization of single-cell RNA-seq data in R. Bioinformatics 33, 1179–1186.

Meulemans, D., and Bronner-Fraser, M. (2004). Gene-regulatory interactions in neural crest evolution and development. Dev Cell 7, 291–299.

Mi, H., Huang, X., Muruganujan, A., Tang, H., Mills, C., Kang, D., and Thomas, P.D. (2017). PANTHER version 11: expanded annotation data from Gene Ontology and Reactome pathways, and data analysis tool enhancements. Nucleic Acids Res 45, D183–D189.

Milo, R., Shen-Orr, S., Itzkovitz, S., Kashtan, N., Chklovskii, D., and Alon, U. (2002). Network motifs: simple building blocks of complex networks. Science 298, 824–827.

Nam, J., and Davidson, E.H. (2012). Barcoded DNA-tag reporters for multiplex cis-regulatory analysis. PLoS One 7, e35934.

Nitzan, E., Krispin, S., Pfaltzgraff, E.R., Klar, A., Labosky, P.A., and Kalcheim, C. (2013). A dynamic code of dorsal neural tube genes regulates the segregation between neurogenic and melanogenic neural crest cells. Development 140, 2269–2279.

Noble, W.S. (2009). How does multiple testing correction work? Nat Biotechnol 27, 1135–1137.

Ochoa, S.D., Salvador, S., and LaBonne, C. (2012). The LIM adaptor protein LMO4 is an essential regulator of neural crest development. Dev Biol 361, 313–325.

Olesnicky Killian, E.C., Birkholz, D.A., and Artinger, K.B. (2009). A role for chemokine signaling in neural crest cell migration and craniofacial development. Dev Biol 333, 161–172.

Osterwalder, M., Barozzi, I., Tissieres, V., Fukuda-Yuzawa, Y., Mannion, B.J., Afzal, S.Y., Lee, E.A., Zhu, Y., Plajzer-Frick, I., Pickle, C.S., et al. (2018). Enhancer redundancy provides phenotypic robustness in mammalian development. Nature 554, 239–243.

Peter, I.S., Faure, E., and Davidson, E.H. (2012). Predictive computation of genomic logic processing functions in embryonic development. Proc Natl Acad Sci U S A 109, 16434–16442.

Peukert, D., Weber, S., Lumsden, A., and Scholpp, S. (2011). Lhx2 and Lhx9 determine neuronal differentiation and compartition in the caudal forebrain by regulating Wnt signaling. PLoS Biol 9, e1001218.

Picelli, S., Faridani, O.R., Bjorklund, A.K., Winberg, G., Sagasser, S., and Sandberg, R. (2014). Full-length RNA-seq from single cells using Smart-seq2. Nat Protoc 9, 171–181.

Preger-Ben Noon, E., Sabaris, G., Ortiz, D.M., Sager, J., Liebowitz, A., Stern, D.L., and Frankel, N. (2018). Comprehensive Analysis of a cis-Regulatory Region Reveals Pleiotropy in Enhancer Function. Cell Rep 22, 3021–3031.

Prescott, S.L., Srinivasan, R., Marchetto, M.C., Grishina, I., Narvaiza, I., Selleri, L., Gage, F.H., Swigut, T., and Wysocka, J. (2015). Enhancer divergence and cis-regulatory evolution in the human and chimp neural crest. Cell 163, 68–83.

Rada-Iglesias, A., Bajpai, R., Prescott, S., Brugmann, S.A., Swigut, T., and Wysocka, J. (2012). Epigenomic annotation of enhancers predicts transcriptional regulators of human neural crest. Cell Stem Cell 11, 633–648.

Ramirez, F., Ryan, D.P., Gruning, B., Bhardwaj, V., Kilpert, F., Richter, A.S., Heyne, S., Dundar, F., and Manke, T. (2016). deepTools2: a next generation web server for deep-sequencing data analysis. Nucleic Acids Res 44, W160–165.

Sandmann, T., Girardot, C., Brehme, M., Tongprasit, W., Stolc, V., and Furlong, E.E. (2007). A core transcriptional network for early mesoderm development in Drosophila melanogaster. Genes Dev 21, 436–449.

Sauka-Spengler, T., and Barembaum, M. (2008). Gain- and loss-of-function approaches in the chick embryo. Methods Cell Biol 87, 237–256.

Sauka-Spengler, T., and Bronner-Fraser, M. (2008). A gene regulatory network orchestrates neural crest formation. Nat Rev Mol Cell Biol 9, 557–568.

Schulz, Y., Wehner, P., Opitz, L., Salinas-Riester, G., Bongers, E.M., van Ravenswaaij-Arts, C.M., Wincent, J., Schoumans, J., Kohlhase, J., Borchers, A., et al. (2014). CHD7, the gene mutated in CHARGE syndrome, regulates genes involved in neural crest cell guidance. Hum Genet 133, 997–1009

Simoes-Costa, M., and Bronner, M.E. (2015). Establishing neural crest identity: a gene regulatory recipe. Development 142, 242–257.

Simoes-Costa, M., and Bronner, M.E. (2016). Reprogramming of avian neural crest axial identity and cell fate. Science 352, 1570–1573.

Simoes-Costa, M., Tan-Cabugao, J., Antoshechkin, I., Sauka-Spengler, T., and Bronner, M.E. (2014). Transcriptome analysis reveals novel players in the cranial neural crest gene regulatory network. Genome Res 24, 281–290.

Simoes-Costa, M.S., McKeown, S.J., Tan-Cabugao, J., Sauka-Spengler, T., and Bronner, M.E. (2012). Dynamic and differential regulation of stem cell factor FoxD3 in the neural crest is Encrypted in the genome. PLoS Genet 8, e1003142.

Skene, P.J., and Henikoff, S. (2017). An efficient targeted nuclease strategy for high-resolution mapping of DNA binding sites. Elife 6.

Smith, C.L., and Tallquist, M.D. (2010). PDGF function in diverse neural crest cell populations. Cell Adh Migr 4, 561–566.

Smith, J., Theodoris, C., and Davidson, E.H. (2007). A gene regulatory network subcircuit drives a dynamic pattern of gene expression. Science 318, 794–797.

Spitz, F., Gonzalez, F., and Duboule, D. (2003). A global control region defines a chromosomal regulatory landscape containing the HoxD cluster. Cell 113, 405–417.

Stark, R.B., G. D. (2011). DiffBind: differential binding analysis of ChIP-seq peak data. Bioconductor http://bioconductor.org/packages/release/bioc/html/DiffBind.html.

Talbot, D., Collis, P., Antoniou, M., Vidal, M., Grosveld, F., and Greaves, D.R. (1989). A dominant control region from the human beta-globin locus conferring integration site-independent gene expression. Nature 338, 352–355.

Trainor, P.A. (2010). Craniofacial birth defects: The role of neural crest cells in the etiology and pathogenesis of Treacher Collins syndrome and the potential for prevention. Am J Med Genet A 152A, 2984–2994.

Trainor, P.A., and Krumlauf, R. (2000). Patterning the cranial neural crest: hindbrain segmentation and Hox gene plasticity. Nat Rev Neurosci 1, 116–124.

Verzi, M.P., Agarwal, P., Brown, C., McCulley, D.J., Schwarz, J.J., and Black, B.L. (2007). The transcription factor MEF2C is required for craniofacial development. Dev Cell 12, 645–652.

Waimey, K.E., Huang, P.H., Chen, M., and Cheng, H.J. (2008). Plexin-A3 and plexin-A4 restrict the migration of sympathetic neurons but not their neural crest precursors. Dev Biol 315, 448–458.

Whyte, W.A., Orlando, D.A., Hnisz, D., Abraham, B.J., Lin, C.Y., Kagey, M.H., Rahl, P.B., Lee, T.I., and Young, R.A. (2013). Master transcription factors and mediator establish super-enhancers at key cell identity genes. Cell 153, 307–319.

Williams, R.M., Senanayake, U., Artibani, M., Taylor, G., Wells, D., Ahmed, A.A., and Sauka-Spengler, T. (2018). Genome and epigenome engineering CRISPR toolkit for in vivo modulation of cis-regulatory interactions and gene expression in the chicken embryo. Development 145.

Wyrick, J.J., and Young, R.A. (2002). Deciphering gene expression regulatory networks. Curr Opin Genet Dev 12, 130–136.

Ye, T., Krebs, A.R., Choukrallah, M.A., Keime, C., Plewniak, F., Davidson, I., and Tora, L. (2011). seqMINER: an integrated ChIP-seq data interpretation platform. Nucleic Acids Res 39, e35.

Zhu, L., Zhang, S., and Jin, Y. (2014). Foxd3 suppresses NFAT-mediated differentiation to maintain self-renewal of embryonic stem cells. EMBO Rep 15, 1286–1296.

Zinzen, R.P., Girardot, C., Gagneur, J., Braun, M., and Furlong, E.E. (2009). Combinatorial binding predicts spatiotemporal cis-regulatory activity. Nature 462, 65–70.

