## Supplemental_Material for "Reconstruction of the global neural crest gene regulatory network *in vivo*"

[1]University of Oxford, MRC Weatherall Institute of Molecular Medicine, Radcliffe Department of Medicine, Oxford, OX3 9DS, UK
[2]University of Oxford, MRC Centre for Computational Biology, MRC Weatherall Institute of Molecular Medicine, Oxford, OX3 9DS, UK
[3]University of Oxford, MRC Molecular Haematology Unit, MRC Weatherall Institute of Molecular Medicine, Oxford, OX3 9DS, UK
[4]Present Address: Okinawa Institute of Science and Technology, Molecular Genetics Unit, Onna, 904-0495, Japan

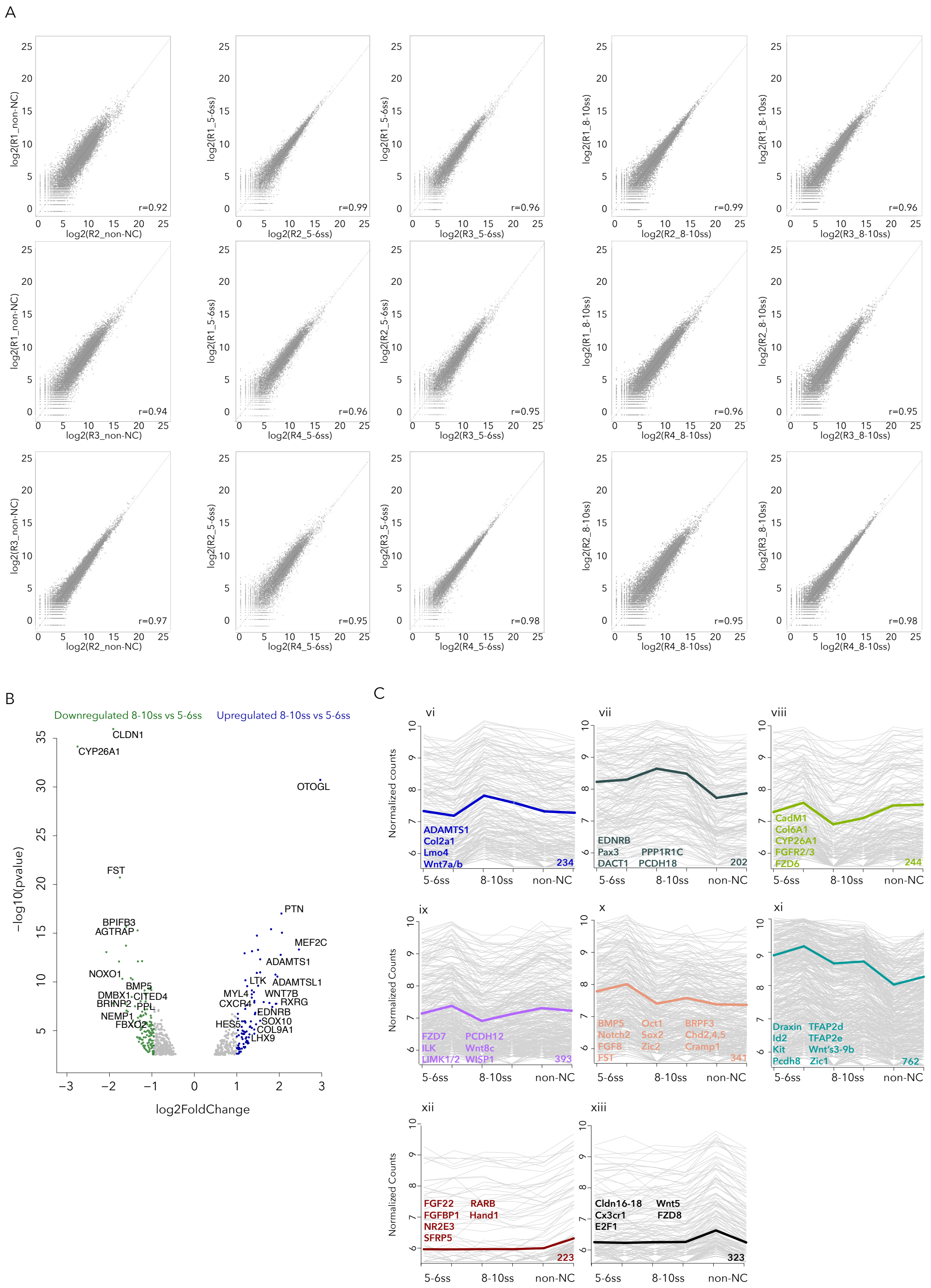

**Supplementary Figure S1. RNA-seq quality control and reproducibility. Related to figure 1 (A)** Scatter plots showing correlation of RNA-seq replicas. r= Pearson correlation co-efficient. **(B)** Volcano plot of genes up and downregulated at 8-10ss compared to 5-6ss. **(C)** Clusters (vi-xiii) of highly correlated genes identified by WGCNA.

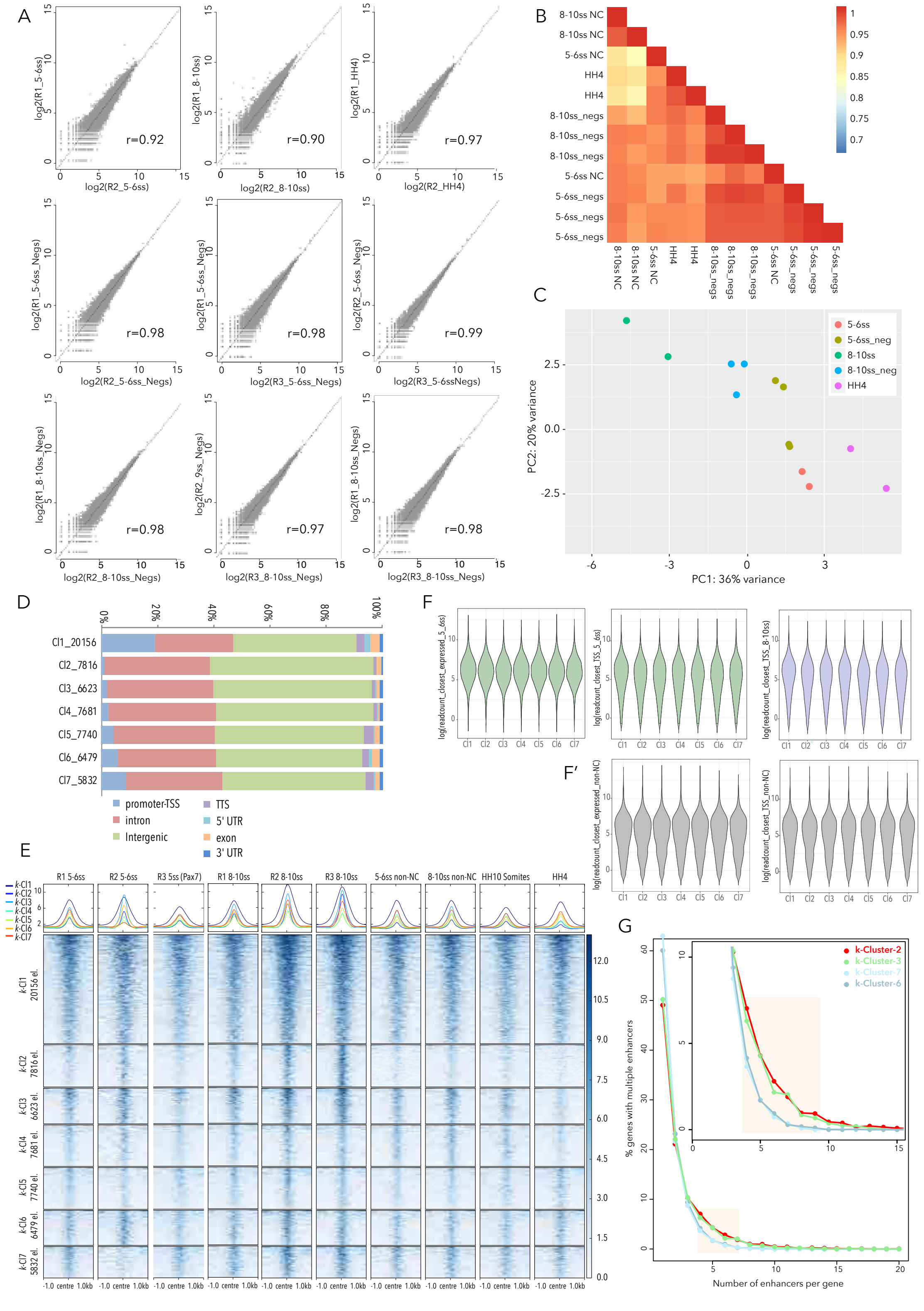

**Supplementary Figure S2. ATAC-seq quality control and chromatin accessibility dynamics. Related to Figures 2 and 3.** **(A)** Scatter plots showing correlation of ATAC-seq replicas, r= Pearson correlation co-efficient. **(B)** Matrix presenting the correlation coefficients to all possible pairwise comparisons of replicates/samples. **(C)** PCA comparing NC and non-NC cells at both stages and HH4 ATAC-seq samples. **(D)** Stacked bar plot showing genomic annotation of *k*-Cluster elements. The number of elements in each *k*-Cluster is also shown. **(E)** Heatmap and merged profiles depicting *k*-means linear enrichment clustering of ATAC signal across all samples/stages analysed. Pax7 sample is NC cells isolated using the Pax7-195 enhancer (Fig. S3). **(F, F′)** Violin plots showing correlation between *k*-Cluster elements and gene expression levels in NC cells (F, green, 5-6ss, purple, 8-10ss) and non-NC cells (F′) annotated to closest expressed gene and closest TSS. **(G)** Percentage of genes with multiple associated enhancers.

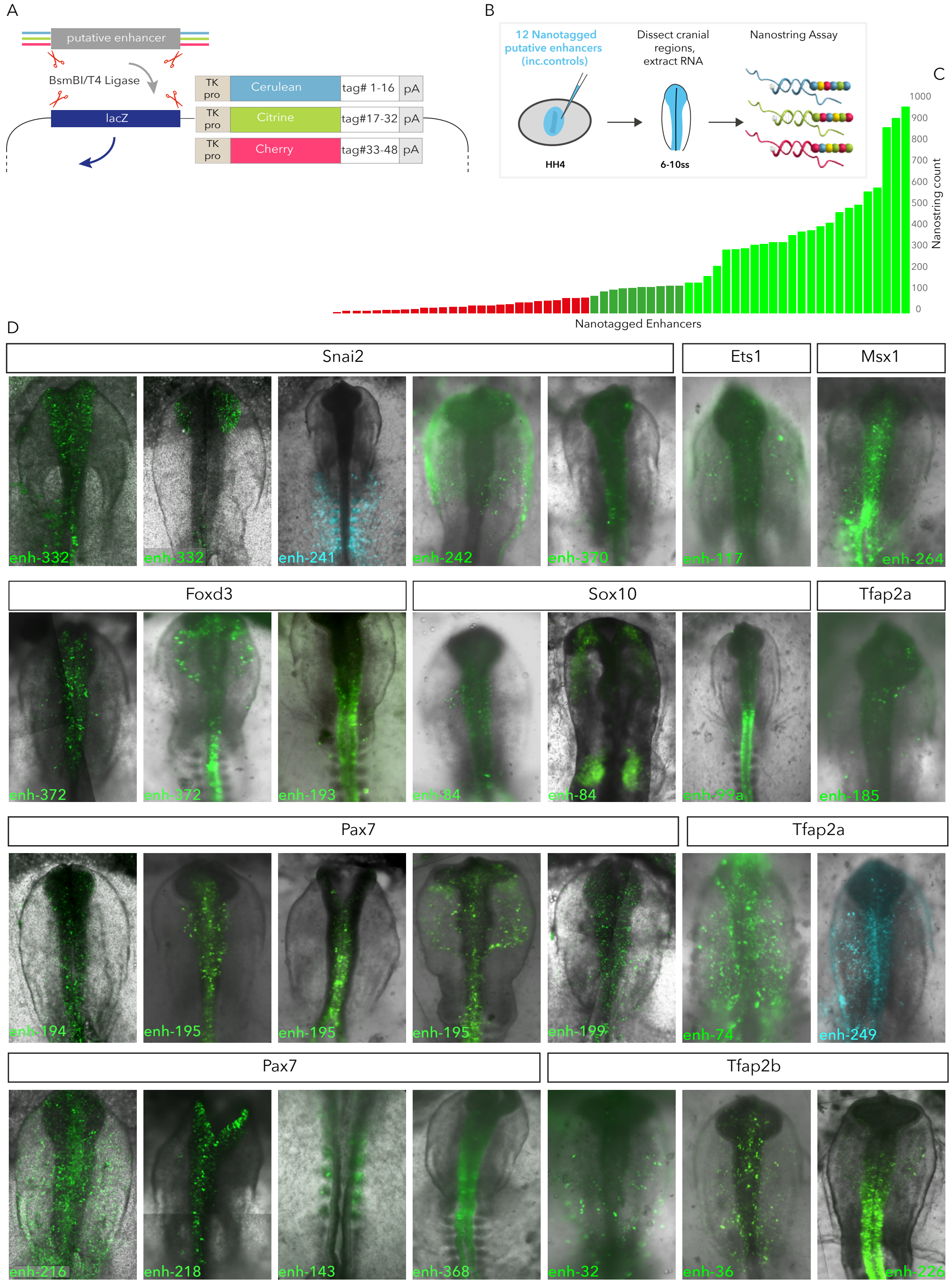

**Supplementary Figure S3. Multiplexed high-throughput enhancer screening. Related to Figures 3 and 4. (A)** Schematic depiction of enhancer cloning strategy. **(B)** Cartoon showing *ex ovo* electroporation technique and Nanostring assay. **(C)** Bar graph representing typical Nanostring results. Nanostring count (of nanotag transcripts) above 50 (green) was determined to reflect *in vivo* enhancer activity. **(D)** *In vivo* activity of selected enhancers.

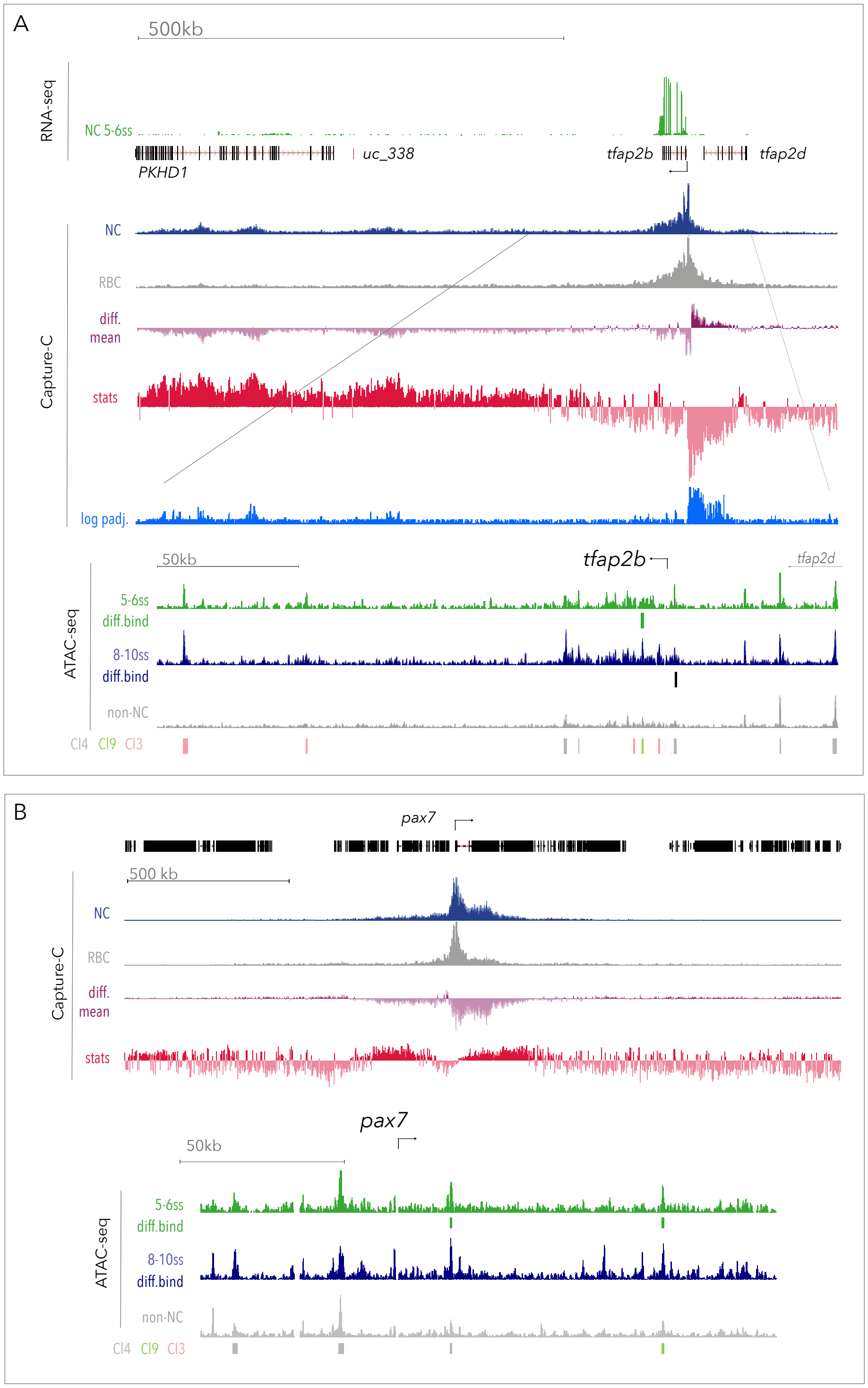

**Supplementary Figure S4. Capture-C at *Tfap2b* and *Pax7* loci. Related to Figure 3.** Genome browser views of chromosome conformation capture (Capture-C) from the *Tfap2b* **(A)** and *Pax7* **(B)** promoters and associated statistical analysis of the differences between Capture-C profiles in neural crest (NC, n=4) and red blood cells (RBC, n=4). Raw counts of unique interactions mapped to each restriction fragment were analysed using the bioconductor package DESeq2. **(A)** shows $\sim$1Mb around the *Tfap2b* locus, the topologically associating domain mostly spanning the region downstream of the *Tfap2b* coding region. **(B)** shows $\sim$2Mb around the *Pax7* locus, the topologically associating domain mostly spanning the gene body and downstream of the *Pax7* coding region. The red line denotes the position of the capture probe. The top track shows the NC RNA-seq output and the second and third tracks show the Capture-C normalised counts from raw count per restriction fragment from NC cells (in blue) and RBCs (in grey), respectively. Purple track shows differential mean between NC and RBC profiles specifically highlighting proximal and distal interaction blocks, with NC-specific interactions overlapping distal *cis*-regulatory elements (ATAC-seq tracks and mapped analysed *k*-Cluster and Diffbind elements). The majority of differences are with elements that interact more strongly in NC than in RBCs and the DESeq2 analysis highlighted these interactions as being statistically significant. Statistical significance is presented in the form of the DEseq2 Wald statistics track (in red), which determines significance of difference in interactions between NC and RBCs, and is calculated as a ratio of LogFoldChange values and their standard errors. Blue track showing the log transformed p-values mapped across the locus indicates the significant differences between NC and RBC profiles, and points to both proximal and distal interactions.

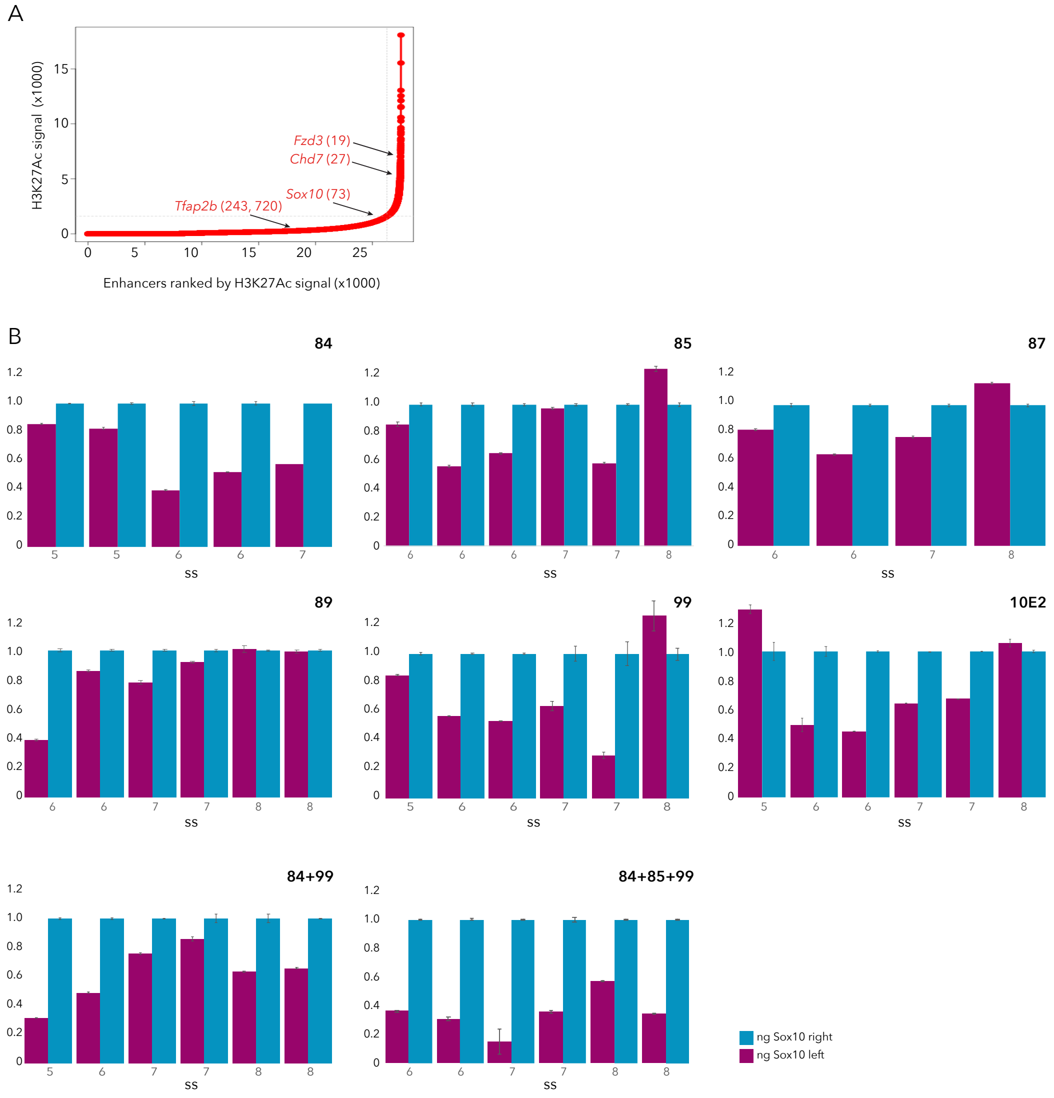

**Supplementary Figure S5. Decommissioning super-enhancer elements controlling *Sox10* expression. Related to Figure 4. (A)** Enhancers ranked by H3K27ac signal from 5-6ss, using the ROSE algorithm, top-ranked genes are annotated. **(B)** qPCR for *Sox10* following dCas9-Krab mediated decommissioning of associated enhancers using bilateral electroporation (Fig. 4K-M). *Sox10* on the left (experimental) side of embryos shown in magenta, right (control) side shown in blue. Error bars show standard deviation.

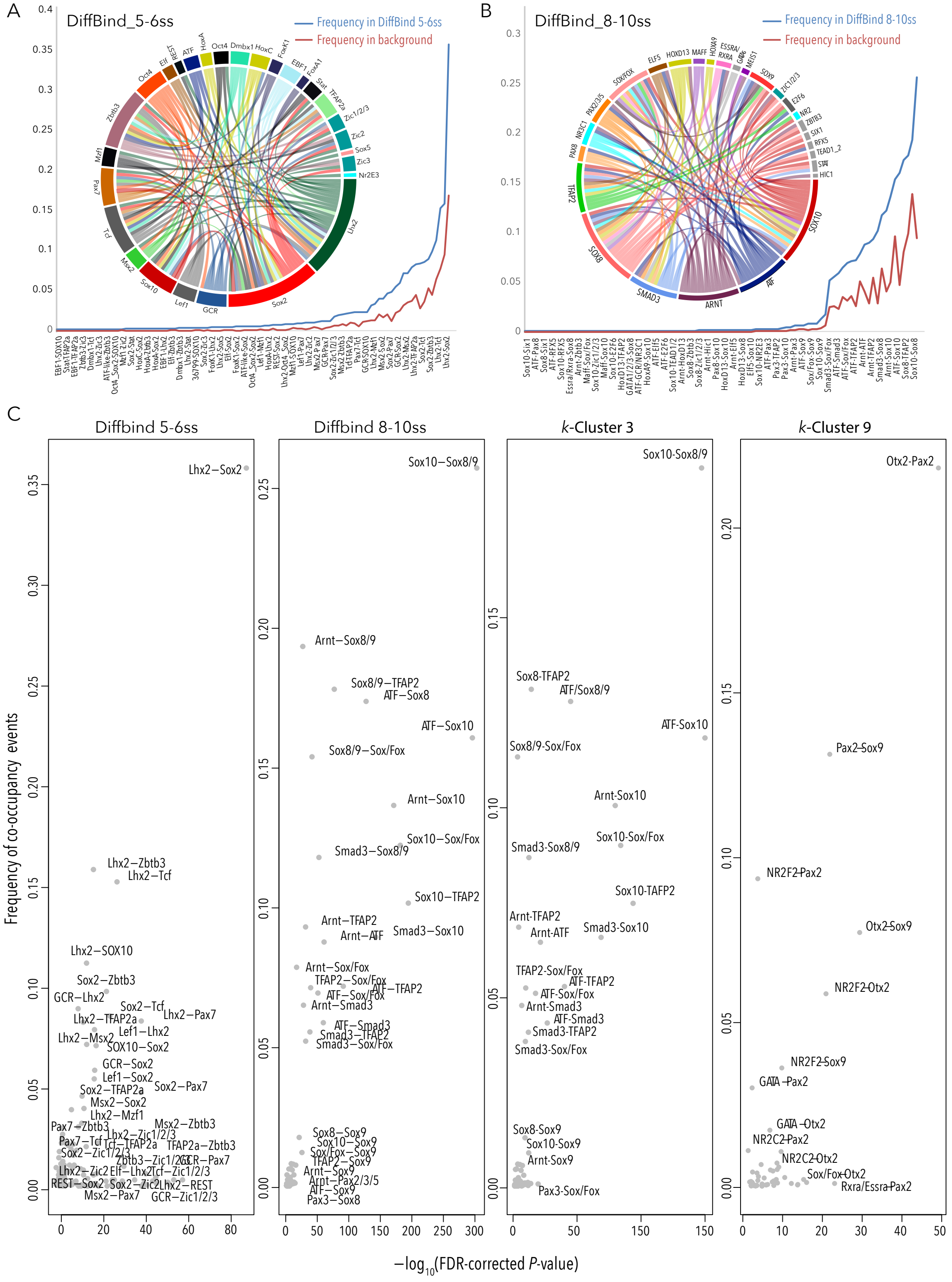

**Supplementary Figure S6. Combinatorial transcription factor binding in DiffBind elements. Related to Figure 6.** **(A, B)** Motif co-occurrence frequencies and circular networks plots showing putative TF combinatorial binding interactions in DiffBind 5-6ss **(A)** and DiffBind 8-10ss **(B)**. **(C)** Showing frequency of TF co-occupancy events and statistical significance in DiffBind 5-6ss, 8-10ss and *k*-Cluster-3 and *k*-Cluster-9

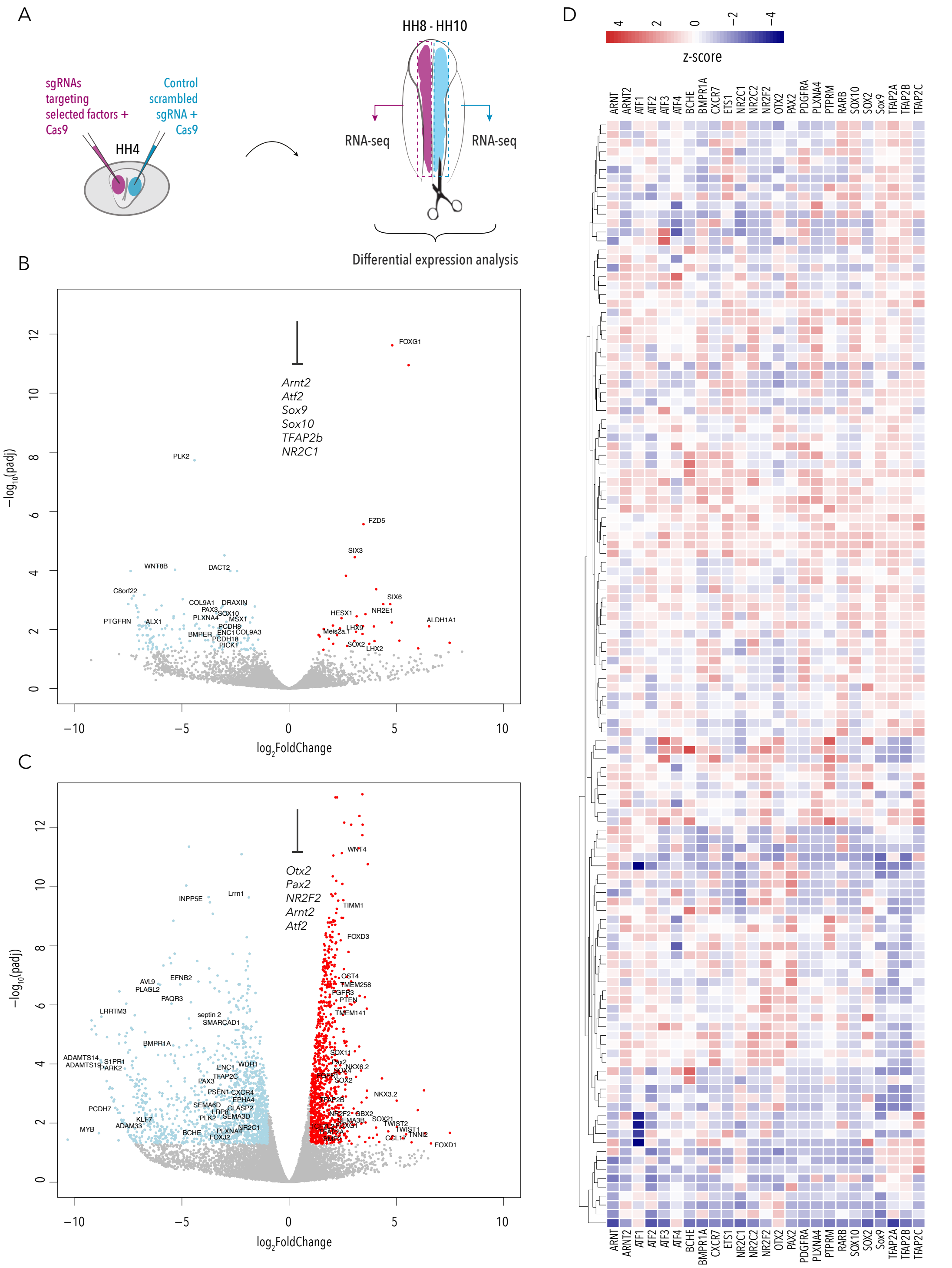

**Supplementary Figure S7. Perturbation of core NC-GRN transcription factors. Related to Figure 6. (A)** Schematic of bilateral electroporation assay for CRISPR mediated perturbation experiments. **(B, C)** Volcano plots showing up- and down-regulated genes following CRISPR knock-out of core TFs associated with *k*-Cluster-3 (B) and *k*-Cluster-9 (C) **(D)** Heatmap showing single-cell co-expression of targeted TFs and selected dysregulated genes following CRISPR knockout of core TFs..

**Table S1.** Capture-C targets and oligo sequences

| | GalGal4 location | Capture oligo sequence |
|---|---|---|
| Sox10 | chr1:50,912,325-50,912,444 | TTTTCAAATCAGGGGACAGTGATGCTGTGGCAGGGACTTACAGAGGTGACTGCAGACAGTGAGAGAGGAGGGGTGCACAGGGCAGGCAGCTCAGGTCCTGGGCTTCTCTTCAAGTTGATC |
| | | GATCAAGGACATGCTGGGGATGTGGATACCACAAGCGTGTTGGTGGTGGCGGGGGTAAGGAAGGGGAGCTCAGCCCTGGGTTGCACAACTTCACATCCCTTCCTGTGACCAGCAACTCCA |
| Sox9 | chr18:9,068,402-9,068,521 | AGCCGGGCTGCGCGCTGGTGGAGACTCCGTCTCTGCCGGCTTTACTTCTTGTTTTTAACCCTTCCCCGCCCCTCAGCCGCCCGGTTGTTTTTTTTCTCTCCGTTTTCTCCTCCCCTGATC |
| | | GATCCGCGGGAACCCCTCCGGCACGCAGCGCACGGACTTCGGCGCCTGGGAAGCCCGAAGCCGTCGCGGGGCGGAGCGAAGAGAAGCGCAACGGCTCCCACCGCCCCGCGCCCCGCCCCG |
| Lmo4 | chr8:14,676,982-14,677,101 | GCTTTTTTAATGGAGACGGAGGGAGGGACGCGCGGAGAGCTGGCAATTTGTAGGACGAAAATGGATGCTTAATTCACGTCTCGGTTTTAATTAGGTGATTCACCGGATTTCTCCCGGATC |
| | | GATCTTCCCCCCGCAGCCGGCGCACCTCTTCCACGAGAGGGAGCCCGCCGTCACCGGGGGCGGCTGCGCGCTGCCGCCGGGGTTCACCATGGCCTGCGGGAGAGCGGCGCGGGTCAGGGG |
| Pax7 | chr21:4,443,114-4,443,233 | TTTGGGGCGGTTGGAGCTCCTTTCCCACGCCGCGCCTTTCCCCGAGCAGCTGTGCCGCTTTGCTCTTTATTTCTCCTCCCGTTTCAAGTAGTGAGGAGCCGGCTTTCAGAAGCCAGGATC |
| | | GATCCAATTCATTAAGGATGCTAATGAAGGAGGTGCGTCGGGAGCCGCCGAGCGCAGAGCGGAGGGGTTTGGGTCTCATTTCGGCCCTATATACGGGGGGGTGATGTGGAAGGAATGCTG |
| Snai2 | chr2:108,126,987-108,127,106 | GATCCTCTTTGAATAACTGAGTTCAAGTGGATGAACAAACTTCATGATTCATTCCGAGCAGCGCTGACATATTTGTCCGAACTGCCTCACTGTAAGCACGGCAAAAAAACAGAGCTAGCG |
| | | GGGAGAAGCGCAGCTCTTCACAGCACTGAGGGCAAAGCTGCTGCTTTCCTTCACTGTACAGAAACGATTAAATCCACTTTTGGAAGGGACGTTCGTGACGCCGGTCGCCTCTCAGCGATC |
| TFAP2b | chr3:107,873,143-107,873,262 | GATCCATCTATAATTGGAAATGGGGGACAGACACCAAATCCGACGTTCCTCTTCCATCGCAACTTATATCTGTTGTCTGAAACAATAGGCTGCAGAAGTAAACCTCAATCGGATAGTAAA |
| | | GACATATATAAAGTGACCCATTTATATATGTAAATTATATATATTCTCGCTTATGTATGGATTTACATAGGCACATGTATGCTACACGTTACATATGCATATATGCATACAAACCTGATC |
| Zfhx4 | chr2:119,024,967-119,025,086 | TTTTGGAACAGCTGTAAATTAGTGATGAGCTATTAGTGAGCTGTGTCATTATTTAATAAAAATGGCTTCTCTCACCTTATTTTTTATCCAGGTCCCTGACAGGCTGGATGAAATGAGATC |
| | | GATCTGCAAGGACTCGGAGTAGCTTAGGCTGTAATCAGGCTACTATTACAGTGGCTGGAGCCTTGCAGGCTCCCAAAAAAATGAAGGAAAGCAGACTTTTAACCAATGTGTGACCAACTT |
| $\alpha$Globin | chr14:12,097,459-12,097,578 | GATCCTAACACTAACCCCAGCTCGCGTCGGGGTCCAACCCCCCCAGCCTGCGCAGTATCGTGGGTGGGGCAGGGCAGCAGCCCTGCCTGGCTGGGGTCCAGAATCTATGGGGCGGGCTGG |
| | | ACAAGAACAACGTCAAGGGCATCTTCACCAAAATCGCCGGCCATGCTGAGGAGTATGGCGCCGAGACCCTGGAAAGGTAGGTGTCCTTCTCTGTCCTCCGGCTGCCTCTCTCCCCTGATC |

**Table S2.** Primer design for mulitplex enhancer cloning and Nanostring screening.

| | | BsmBI binding site | Vector specific overhangs | Target specific seqeunce ~20nt |
|---|---|---|---|---|
| Cerulean vectors F | TTTTTT | CGTCTC | ccatgg | nnnnnnnnnnnnnnnnnnn |
| Cerulean vectors R | TTTTTT | CGTCTC | ggtcct | nnnnnnnnnnnnnnnnnnn |
| Citrine vectors F | TTTTTT | CGTCTC | gccagg | nnnnnnnnnnnnnnnnnnn |
| Citrine vectors R | TTTTTT | CGTCTC | caacag | nnnnnnnnnnnnnnnnnnn |
| Cherry vectors F | TTTTTT | CGTCTC | gtgcag | nnnnnnnnnnnnnnnnnnn |
| Cherry vectors R | TTTTTT | CGTCTC | caccgt | nnnnnnnnnnnnnnnnnnn |
| Cerulean neg control oligo F | TTTTTT | CGTCTC | ccatgg | AGCTGGATCGATgatatcCGATCGATCGTAGCAC |
| Cerulean neg control oligo R | TTTTTT | CGTCTC | ggtcct | GTGCTACGATCGATCGgatatcATCGATCCAGCT |
| Citrine neg control oligo F | TTTTTT | CGTCTC | gccagg | AGCTGGATCGATgatatcCGATCGATCGTAGCAC |
| Citrine neg control oligo R | TTTTTT | CGTCTC | caacag | GTGCTACGATCGATCGgatatcATCGATCCAGCT |
| Cherry neg control oligo F | TTTTTT | CGTCTC | gtgcag | AGCTGGATCGATgatatcCGATCGATCGTAGCAC |
| Cherry neg control oligo R | TTTTTT | CGTCTC | caccgt | GTGCTACGATCGATCGgatatcATCGATCCAGCT |

**Table S3**. Guide RNA sequences

| Epigenome engineering of *Sox10* enhancers | | | Targeted knock-out of core TFs | |
|---|---|---|---|---|
| Target/sgRNA | sgRNA seqeunce | | Target/sgRNA | sgRNA seqeunce |
| 84_sgRNA_1 | AGTCTGCCACCCATCAAAGC | | ATF2_sgRNA_1 | TCAACAACTGAAACACCGgt |
| 84_sgRNA_2 | CCATTGTATCATGCTGGACA | | ATF2_sgRNA_2 | cttgctgttttcagGCATCA |
| 84_sgRNA_3 | CTCCACTGAACGAGTCCATG | | | |
| 84_sgRNA_4 | ATTAATTCCTGCGAACAGAA | | TFAp2b_sgRNA_1 | GGAGGAGTGCTGAGAAGgta |
| 84_sgRNA_5 | CCCTTTGTGTATGGGCTCAC | | TFAp2b_sgRNA_2 | accctcgcttacCTTCCACC |
| 85_sgRNA_1 | AGATGTGCTTATGGGCTCCT | | | |
| 85_sgRNA_2 | TGGGAACAATGTCAACTCCG | | Sox10_sgRNA_1 | ttccctccccagTGAGAAGA |
| 85_sgRNA_3 | GCACAGAGCGGCCCCGTCG | | Sox10_sgRNA_2 | ggtaggaaaacttacATTGC |
| 85_sgRNA_4 | TTCAGTACAGCTACTTACAG | | | |
| 85_sgRNA_5 | TCTTTCCACCCGCCCAGGGC | | Arnt2_sgRNA_1 | AGGGACCCAGCAAATTTTCA |
| 87_sgRNA_1 | CAGGAAGAAATGCGTAGTGA | | Arnt2_sgRNA_2 | tcttttgtttatagGTATGA |
| 87_sgRNA_2 | GAGCGAGCAGAGAGTGGAGC | | | |
| 87_sgRNA_3 | TCTTTGTTCCCTGCCTTTAA | | NR2C1_sgRNA_1 | CTCTTTACCGCAGCGTATAC |
| 87_sgRNA_4 | CTCTAAAACACCCGATTGTC | | NR2C1_sgRNA_2 | AGACAACTCTCCCAATGAGC |
| 87_sgRNA_5 | GCAGGGAAGGAGGATTCTGA | | | |
| 89_sgRNA_1 | AGGGCATCCCCATGCACAAC | | Sox9_sgRNA_1 | CTCTCATTCAGCAGCCTG |
| 89_sgRNA_2 | GAGGCAACAAATCTTTTCCA | | | |
| 89_sgRNA_3 | AGGCAACTCACTGAGCATGA | | | |
| 89_sgRNA_4 | GGGGAGAGTAAATGAGACAG | | | |
| 89_sgRNA_5 | CAGTCAGTTGGGCTGCAGAG | | | |
| 99_sgRNA_1 | GGTGAGAAATGTTGAAAACG | | | |
| 99_sgRNA_2 | GTGTGTGACTCTTTTGTTCC | | | |
| 99_sgRNA_3 | TGGACAACTTTGCTAGGCCC | | | |
| 99_sgRNA_4 | GCACAAAGGAAATGGATAAA | | | |
| 99_sgRNA_5 | AGGAAAGAGGGAAGGAAGGC | | | |
| 10E2_sgRNA_1 | AGCAGGAGCAGGGAAACAAT | | | |
| 10E2_sgRNA_2 | AAACATAAGCACAAACTAGG | | | |
| 10E2_sgRNA_3 | TGGTAAGGATGGCCTGGATC | | | |
| 10E2_sgRNA_4 | GCTGGGGAGGGGAGGCGGGC | | | |
| 10E2_sgRNA_5 | CCATATCAACCATTCTCCAG | | | |
| Scrambled control | TGCAGTGCTTCAGCCGCT | | | |
